# An updated antennal lobe atlas for the yellow fever mosquito *Aedes aegypti*

**DOI:** 10.1101/865675

**Authors:** Shruti Shankar, Conor J. McMeniman

## Abstract

The yellow fever mosquito *Aedes aegypti* is a prolific vector of arboviral and filarial diseases that largely relies on its sense of smell to find humans. To facilitate in-depth analysis of the neural circuitry underlying *Ae. aegypti* olfactory-driven behaviors, we generated an updated *in vitro* atlas for the antennal lobe olfactory brain region of this disease vector using two independent neuronal staining methods. We performed morphological reconstructions with replicate fixed, dissected and stained brain samples from adult male and female *Ae. aegypti* of the LVPib12 genome reference strain and determined that the antennal lobe in both sexes is comprised of approximately 80 discrete glomeruli. Guided by landmark features in the antennal lobe, we found 63 of these glomeruli are stereotypically located in spatially invariant positions within these *in vitro* preparations. A posteriorly positioned, mediodorsal glomerulus denoted MD1 was identified as the largest spatially invariant glomerulus in the antennal lobe. Spatial organization of glomeruli in a recently field-derived strain of *Ae. aegypti* from Puerto Rico was conserved, despite differences in antennal lobe shape relative to the inbred LVPib12 strain. This model *in vitro* atlas will serve as a useful community guide and resource to improve antennal lobe annotation and anatomically map projection patterns of neurons expressing target genes in this olfactory center. It will also facilitate the development of chemotopic maps of odor representation in the mosquito antennal lobe to decode the molecular and cellular basis of *Ae. aegypti* attraction to human scent and other chemosensory cues.

**Author Summary:** The olfactory system of the yellow fever mosquito *Aedes aegypti* is highly tuned for the detection of human odorants, as well as other chemical cues influencing host and food-search behavior, egg-laying and mating. To provide insights into the neuroanatomical organization of the olfactory system of this globally important disease vector, we have generated an updated *in vitro* atlas for the primary smell processing center of the *Ae. aegypti* brain, called the antennal lobe. These new guide maps facilitate systematic interrogation of antennal lobe morphology and naming of associated substructures in dissected brain samples of this species labeled with two common neural staining methods. We report that landmark features of the *Ae. aegypti* antennal lobe morphology and spatial organization appear conserved between mosquito sexes and across geographically divergent strains of this mosquito species. An improved understanding of *Ae. aegypti* antennal lobe neuroanatomy and how attractive or repellent odorant stimuli are encoded in this brain center has the potential to rapidly accelerate reverse engineering of synthetic chemical blends that effectively lure, confuse or repel this major disease vector.

## Introduction

*Aedes aegypti* is a prolific vector of yellow fever, dengue, chikungunya, Zika and filariasis in tropical and subtropical regions around the world. This highly anthropophilic mosquito species uses its sense of smell to track volatile chemicals present in human skin odor and breath and orientate towards us. For example, carbon dioxide (1) and other volatile constituents of human body odor, including lactic acid (2), trigger behavioral activation and steer directional flight of this mosquito species towards humans to acquire a blood (3). Olfaction also influences other critical aspects of *Ae. aegypti* biology including oviposition (4), mating (5) and nectar-seeking (6). Molecular and cellular substrates underlying the *Ae. aegypti* sense of smell therefore represent key targets for the control of mosquito-borne diseases transmitted by this cosmopolitan and urbanized disease vector.

In the *Ae. aegypti* adult, olfactory sensory neurons (OSNs) expressing chemoreceptors that detect odorants are primarily localized to peripheral olfactory organs (1, 7–10) including the antennae, maxillary palps and proboscis. Dendritic sensory nerve endings of OSNs are found in various morphological classes of specialized porous sensilla distributed along the length of these organs. Each *Ae. aegypti* sensillum typically houses 2-3 OSNs (7–9). The axonal processes of these neurons project centrally to the mosquito brain, to the first center of olfactory information processing called the antennal lobes. The antennal lobes are made up of clusters or foci of synaptic connectivity called glomeruli, where olfactory information is transmitted from OSNs to second order neurons (11–13).

In the vinegar fly *Drosophila melanogaster*, an individual glomerulus is the point at which the axonal projections of OSNs typically expressing a unique complement of ligand-binding chemoreceptors converge (13, 14) and synapse with the dendritic arbors of projection neurons (PNs) and local interneurons (LNs) (15). Each glomerulus is therefore tuned to a unique subset of chemicals. Anterograde dye-filling of peripheral OSNs that respond functionally to the same subset of odors and subsequent tracing of their glomerular projections indicates this pattern is likely to be conserved in mosquitoes (12, 16, 17).

An initial morphological study of the *Ae. aegypti* antennal lobe by Bausenwein and Nick (18) denoted the presence of approximately 35 antennal lobe glomeruli in a single *Ae. aegypti* female examined. However, it was noted that due to the staining methods employed in this study that subdivision of antennal lobe glomerular neuropil in some areas was unclear. Subsequently, Ignell and coauthors (11) estimated that the antennal lobes of this species were comprised of between 49-50 glomeruli using improved immunohistochemical techniques.

A significant volume of the *Ae. aegypti* antennal lobes was previously ascribed a non-olfactory mechanosensory function by Ignell (11). This large core of neuropil on the anterior surface of the antennal lobe was annotated as the Johnston’s Organ Centre (JOC) based on silver nitrate Golgi staining. This analysis revealed the putative sensory innervation of this antennal lobe region by afferent fibers from the Johnston’s Organ (JO), an observation mirrored shortly thereafter by the same authors in *Anopheles gambiae* (19). However, a recent reassessment of antennal lobe morphology in *Anopheles coluzzii* failed to observe the JOC in the antennal lobes of this species, instead identifying discrete glomeruli in this spatial region (20).

To assist basic and applied studies of *Ae. aegypti* olfaction, we have used improved neuropil labeling and confocal imaging techniques to re-examine the neuroanatomy of the *Ae. aegypti* antennal lobe. We present three-dimensional (3D) models based on manual segmentation and reconstructions of antennal lobe glomeruli from replicate male and female brains from the LVPib12 *Ae. aegypti* genome reference strain. To facilitate spatial orientation and stereotypical identification of glomeruli within the antennal lobe, we also present detailed two-dimensional (2D) maps wherein individual glomeruli are named based on their spatial arrangement in this brain region using nc82- and phalloidin-based neuronal staining methods. Finally, we examine antennal lobe neuroanatomy and organization in a recently field-collected strain of *Ae. aegypti* from the municipality of Patillas, Puerto Rico. Altogether, this work describes a standardized framework for glomerular nomenclature in the *Ae. aegypti* antennal lobe. This atlas will serve as a useful reference for neurogenetic and functional imaging studies of the antennal lobe in this anthropophilic disease vector.

## Results

### Three-dimensional reconstruction of the antennal lobes of male and female *Aedes aegypti* mosquitoes

To define boundaries of the antennal lobe of the LVPib12 genome reference strain of *Ae. aegypti* (21), we initially stained whole dissected and fixed mosquito brains using the monoclonal antibody nc82 as a synaptic marker and acquired 1 µm confocal sections of the central brain starting from the surface of the antennal lobes through the central complex. Similar to other dipteran insects such as *Drosophila melanogaster* (22), the deutocerebrum (11) in *Ae. aegypti* is composed of two paired neuropils: the antennal lobe olfactory center and the periesophageal neuropils which include the antennal mechanosensory and motor center (AMMC).

The left antennal lobe (LAL) and the right antennal lobe (RAL) flank the esophageal foramen and are positioned dorsal to the AMMC (Figures 1a and 1b). The antennal lobes are innervated by the antennal (flagellar) (8) nerves that collectively include the axons of OSNs originating from the third antennal segment (11, 16). In addition, maxillary afferents from the distal segment of the maxillary palp terminate in the antennal lobes (16), supporting its role as the first center in the central mosquito brain to receive signals from the peripheral olfactory organs.

**Figure 1.**
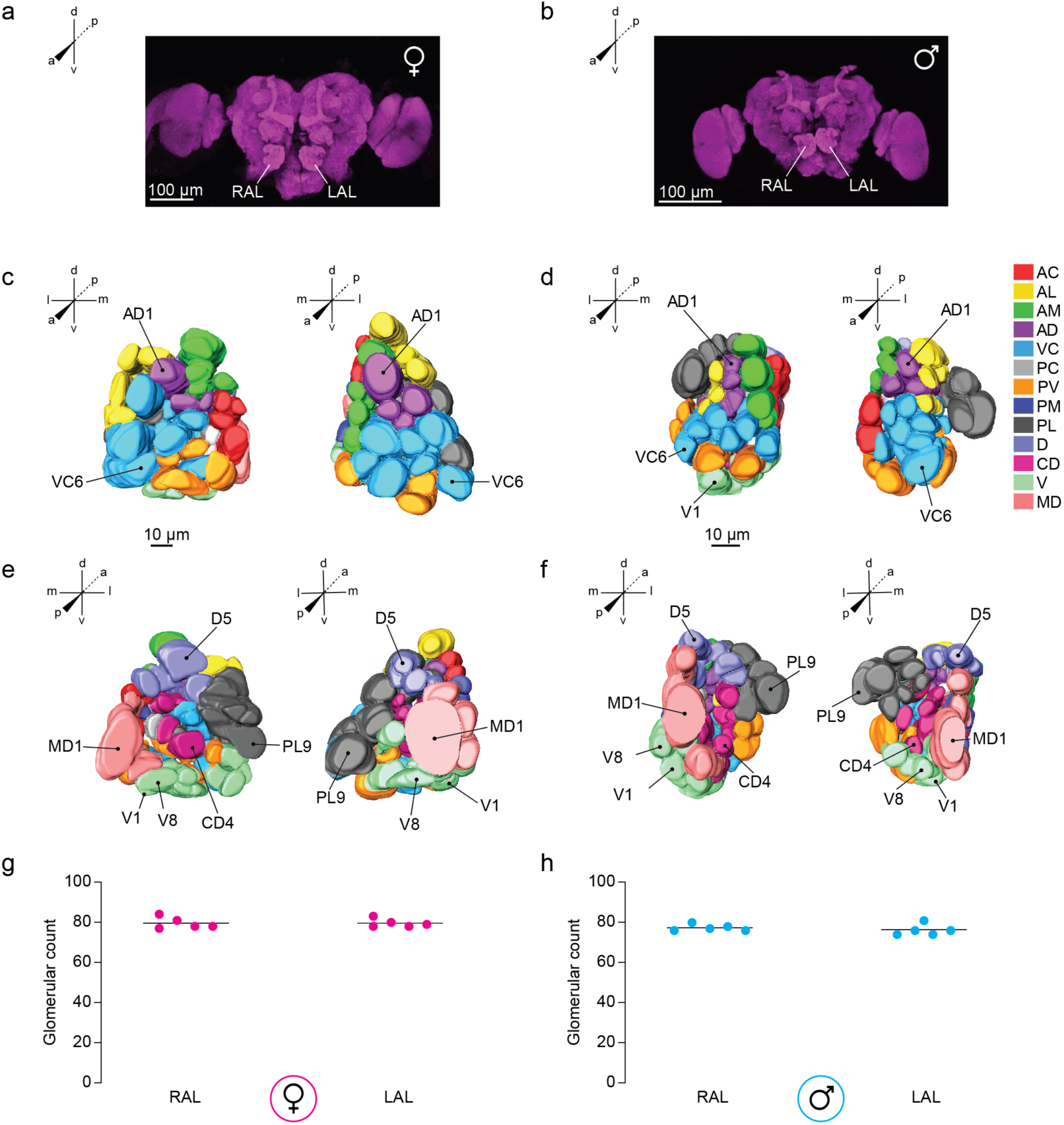
Antennal lobe morphology of the LVPib12 *Aedes aegypti* strain as shown using nc82 staining. Confocal images of a representative adult (**a**) female and (**b**) male brain by nc82 antibody labeling (magenta); Scale bars, 100 µm. Right (RAL) and left antennal lobes (LAL) are indicated. Anterior and posterior views of three-dimensional reconstructive models of the right and left antennal lobe from a female (**c** & **e**) and male (**d** & **f**) mosquito are shown, respectively. Visible landmark glomeruli on these surfaces are denoted. Antennal lobe glomeruli are color-coded according to their classification into 13 groups based on their spatial position along anterior-posterior (a-p), dorsal-ventral (d-v), and medial-lateral (m-l) co-ordinate axes. The total number of glomeruli found in reconstructed (**g**) female and (**h**) male antennal lobes are shown (*n* = 5 brains per sex).

Confocal sections of the LVPib12 antennal lobe revealed that its constituent glomeruli varied both in terms of morphology and volume, and are arranged at different depths relative to i) the anterior to posterior axis of the mosquito body and ii) along the dorsal to ventral neuraxis (22). To re-ascertain glomerular counts relative to previously published studies, we initially reconstructed the left and right antennal lobes from 5 male and 5 female LVPib12 brains that had been stained with nc82 antibodies.

We used the software program Amira to perform segmentation of the antennal lobe by manually painting over all the voxels corresponding to neuropil representing individual glomeruli, which were identified based on their size, shape, spatial position and depth. We found that each antennal lobe in all LVPib12 brains examined was comprised entirely of discrete demarcated glomeruli and we were unable to identify the wedge shaped, JOC non-glomerular core which was described previously (11). The number of glomeruli were found to vary slightly across our reconstructions from 80 ± 2 in the female antennal lobe to 77 ± 2 glomeruli in the male antennal lobe (mean ± s.d.) (Figures 1g and 1h).

### Spatial arrangement of glomeruli in the *Aedes aegypti* antennal lobe

We further sought to understand the pattern of glomerular arrangement within the LVPib12 antennal lobe. Based on spatial position along the anterior-posterior (A-P), ventral-dorsal (V-D) and medial-lateral (M-L) axes, we initially aimed to categorize glomeruli in the male and female antennal lobe into spatial groups, with reference to a previous antennal lobe model (11) that annotated 13 spatial groups of glomeruli in the *Ae. aegypti* antennal lobe. However, as we identified approximately 30 additional glomeruli in our reconstructions relative to this extant map, we first re-annotated spatial descriptors for each group in our atlas to better reflect their spatial positions along these coordinate axes of the antennal lobe (Table S1). These revised groupings organize the *Ae. aegypti* antennal lobe into 13 new glomerular clusters that aid in identifying and naming all glomeruli in each lobe. Each of these groups was assigned a specific color in our reconstructed 3D antennal lobe models (Figure 1c to f).

**Table 1:**
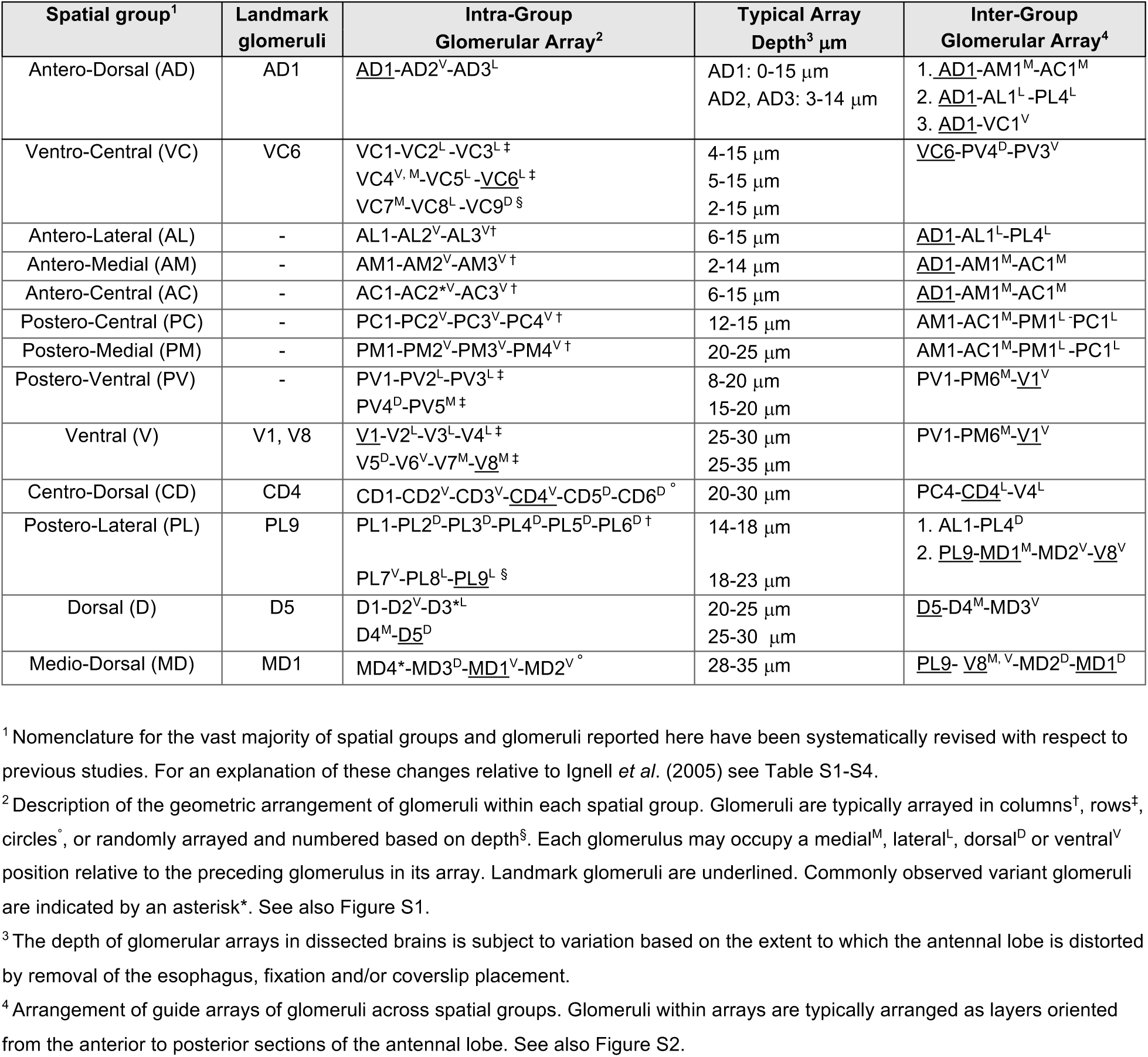
Reference key describing the spatial arrangement of glomeruli in the *Aedes aegypti* antennal lobe.

Viewing the antennal lobe along the A-P axis, on the anterior slices of the antennal lobe we annotated the Antero-Dorsal (AD) group surrounded by the Antero-Lateral (AL) group, laterally; the Antero-Medial (AM) and Antero-Central (AC) groups, medially; and the Ventro-Central (VC) group, ventrally. The groups moving towards the posterior sections of the antennal lobe include the Postero-Medial (PM) group, Postero-Central (PC) and the deeper Medio-Dorsal (MD) groups, along the medial axis; Postero-Lateral (PL) group located on the lateral side; Centro-Dorsal (CD) group located centrally; and Postero-Ventral (PV) group located ventrally. In the most posterior slices, the dorsal group is named Dorsal (D); and the ventrally located group is named Ventral (V).

### A two-dimensional neuroanatomical map of the *Aedes aegypti* antennal lobe

Based on each reconstructed three-dimensional LVPib12 antennal lobe model, we generated two-dimensional maps, showing all manually segmented glomeruli in each confocal slice as observed along the anterior to posterior axis of *in vitro* antennal lobe imaging. As a visual guide, we first selected eight glomeruli with salient features such as a large volume, shape and consistent position across all reconstructions to serve as landmarks for identifying the positions of other glomerular groups. These eight landmark glomeruli are shown in the 3D models (Figure 1c to 1f) and include AD1 and VC6 in anterior antennal lobe slices and D5, PL9, CD4, V1, V8 and MD1 in the more posterior antennal lobe. We did not select landmark glomeruli from the PC, PM, AM and AC groups, as the boundaries of these peripheral glomeruli cannot always be reliably demarcated by the *in vitro* staining methods used here.

To facilitate stereotypical identification of glomeruli based on their relative position to the eight landmark glomeruli, we generated a reference key detailing the typical spatial arrangement of glomeruli in the *Ae. aegypti* antennal lobe (Table 1). In each reconstructed antennal lobe, we assigned a name to each glomerulus based on its spatial group and depth. Glomeruli within the spatial groups are typically arrayed in (i) rows and are numbered based on depth along the medial lateral axis, (ii) in a vertical line (column) or (iii) in a circular manner and numbered along the dorsal-ventral axis. Glomeruli within some spatial groups may also appear to be randomly arrayed. Useful reference arrays of glomeruli within and between spatial groups that assist with identification and naming of glomeruli based on their spatial position relative to landmark glomeruli are detailed in Table 1 and overlaid upon an example antennal lobe reconstruction in Figure S1 (a-j) and Figure S2 (a-m).

Using our reference key, we classified antennal lobe glomeruli as spatially ‘invariant’ or ‘variant’ based on the frequency of identification of individual glomeruli across all reconstructions. Invariant glomeruli were defined as glomeruli with the same assigned name found in the same spatial position in the left and right lobes of both sexes, at a frequency of ≥80%. Variant glomeruli were classified as glomeruli with the same assigned name found with a frequency of <80% across reconstructions, according to these criteria. The number of spatially invariant and variant glomeruli found in each of the 13 spatial groups are described as follows;

The anterior surface of the antennal lobe is comprised of five groups of glomeruli:

1. **Antero-Dorsal group**. In anterior slices of the antennal lobe, AD1 is a landmark glomerulus, distinguishable by its large size and dorsal position. It is located towards the center of the antennal lobe. AD1 is the first antennal lobe glomerulus visualized when stepping down a z-series along the A-P axis, and the typical depth of this glomerulus from its anterior surface (denoted 0 µm) to posterior surface is 15 µm. AM1-3, AC1, VC1, AL1-3 glomeruli routinely cluster around the AD group. AD2 and AD3 are found ventral and towards the medial and lateral side of AD1, respectively. AD2 and AD3 glomeruli are typically found in the space between the AD1 and the VC group and are characteristic satellite glomeruli flanking the AD1 landmark. Three residual variant glomeruli (AD 4-6) are sometimes observed in more dorsal and posterior slices relative to the AD invariant glomeruli. See Figure S1 (a) and Figure S2 (a-e).
2. **Ventro-Central group**: The VC group of glomeruli are found ventral to the AD group in the anterior most slices of the antennal lobe. Previously, a large part of the central antennal lobe was considered to comprise of the Johnston’s Organ Center (JOC). We note that in our reconstructions, this region consists entirely of discrete glomeruli that we have reclassified as the VC and PV spatial groups. The VC group is comprised of an anterior surface layer consisting of the VC1, VC2 and VC3 glomeruli arranged in a row ordered along the medial to lateral axis, or alternatively in a triangle with VC1 positioned dorsal to VC2 and VC3; and the VC4, VC5 and VC6 glomeruli in a more posterior bottom row ordered along the medial to lateral axis. The most posterior layer of this group comprises VC7, VC8 and VC9 that are arranged centrally in this region of the antennal lobe (Table 1). VC1 is a large glomerulus positioned ventral to the landmark glomerulus AD1. VC6 is a large landmark glomerulus located on the lateral side of the antennal lobe and ventral to VC3. Two variant glomeruli (VC10-VC11) in a ventral to dorsal column are sometimes observed more posterior to the VC7-9 cluster. See Figure S1 (b) and Figure S2 (e).
3. **Antero-Lateral group**: The AL group is observed in anterior slices within the dorsal and lateral region of the antennal lobe. The AL1 glomerulus is located laterally, relative to the landmark AD1. AL1, AL2, AL3 glomeruli are positioned in a D-V column on the lateral side of the antennal lobe in order of increasing depth along the A-P axis. For example, AL1 is found dorsal and anterior to AL2. In some reconstructions, three variant glomeruli were sometimes found ventral and posterior to AL3 and were named AL4-AL6. See Figure S1 (c) and Figure S2 (a-b).
4. **Antero-Medial group**: The AM group is located medial to the AD group. It is comprised of the AM1, AM2 and AM3 glomeruli arranged in a column along the D-V axis. AM1 is a large glomerulus located above AD1 towards the medial axis of the antennal lobe. A variant glomerulus AM4 was observed ventral to AM3 in some reconstructions. See Figure S1 (d) and Figure S2 (c-d).
5. **Antero-Central group**: The AC group is located medial to the AM group bisecting the equator of the antennal lobe in a D-V column flanking the medial side. In more posterior slices, the boundaries of some the AC glomeruli were difficult to demarcate as this spatial group is sandwiched between the AM and the more posterior PM spatial groups. AC1 and 3 were spatially invariant glomeruli, whereas variant glomeruli AC2, AC4 and AC5 were observed in some reconstructions. See Figure S1 (e) and Figure S2 (c-d). In mid-sections of the antennal lobe, we describe the following groups:
6. **Postero-Medial group**: This group is located posterior to the AM glomeruli. It is comprised of glomeruli along a D-V column on the medial side of the antennal lobe. The boundaries of PM1, PM2, and PM3 are noted for their small size, and position between the anterior AC and posterior D groups. PM4 is positioned more ventrally and is larger in volume relative to PM1-PM3. In more posterior slices, the ventral PM5 and PM6 are classified as variant glomeruli. See Figure S1 (f) and Figure S2 (d and j).
7. **Postero-Central group**: Included are a small number of glomeruli typically located in between the PM and CD groups. PC1, PC2, PC3 and PC4 glomeruli are arranged in a D-V column. PC1 is found posterior to AC1. In the anterior to posterior direction, AC1-PC1-PM1 glomeruli form an array. A variant glomerulus PC5 can be observed at times ventral to PC4 bordering the ventral group. See Figure S1 (g).
8. **Postero-Ventral group**: The PV group is visible from the anterior to central slices of the antennal lobe. PV glomeruli are found ventral to the VC group. The 5 glomeruli that comprise this group are arranged in two rows ordered along the medial-lateral axis. The ventral row consists of PV1, PV2 and PV3 glomeruli. PV1 flanks the medial antennal lobe margin, with PV2 in the center and PV3 laterally. The dorsal row consists of PV4 and the more posterior PV5. In more posterior slices, variant glomeruli PV6-PV9 may be present. This group of glomeruli were previously considered to be part of the JOC (11), along with the VC spatial group. See Figure S1 (h) and Figure S2 (f and g). The posterior region (−15 µm to −35 µm) of the antennal lobe is comprised of 5 groups of glomeruli. The CD group is located centrally within posterior slices, and the D and V groups are located dorsal and ventral to the CD group respectively. The PL and the MD groups flank the CD group along the lateral and the medial margins of the antennal lobe.
9. **Ventral group**: The V group is found posterior and dorsal to the PV group, in the deepest layers of the antennal lobe. The landmark glomerulus V1 is found ventral and posterior to PV1. V1 flanks the medial margin of the antennal lobe. In the medial to lateral direction the other glomeruli of this group are arranged in three layers. The first layer consists of V1, V2, V3, and V4 aligned in a medial-lateral row. The second more posterior layer consists of V5, V6, V7 and V8 are arranged in an approximate lateral-medial row. V8, which is posterior and medial relative to V7, is a landmark glomerulus that is easy to identify because of its lateral adjacent position to MD1. V8 is found posterior to V1. In some reconstructions we identified more posterior variant V glomeruli (V9-V11). See Figure S1 (i and j) and Figure S2 (g, h and j).
10. **Centro-Dorsal group**: The CD group is the central group in dorsal regions of the posterior-most slices of the antennal lobe. CD1, CD2, CD3, CD4, CD5, and CD6 are arranged almost in a circle. CD1 and CD2 lie centrally in a dorsal position relative to other CD glomeruli, yet ventral to the D group. CD3 and CD4 are more medial in position and ventral to CD1 and CD2. CD5 and the more lateral CD6 are aligned towards the lateral side of the antennal lobe. CD4 is a landmark, conspicuous because it is the most ventral CD glomerulus and due to its large, irregular shape. This glomerulus is located dorsal to the V group. Overall, the CD glomeruli can be distinguished from the other posterior antennal lobe groups by their smaller volume. In some reconstructions, more than 6 CD glomeruli were observed. These variant glomeruli were found in more posterior slices and named CD7-CD12. See Figure S1 (j) and Figure S2 (h).
11. **Postero-Lateral group**: The PL group of glomeruli are located along the lateral axis and form a distinct cluster on the dorso-lateral corner as confocal slices approach the posterior surface of the antennal lobe. This is especially apparent in the male antennal lobe. The PL group lies more posterior than the AL group in both sexes. PL1 is the most ventral glomerulus in this group. Thereafter PL glomeruli are named with increasing numbers based on dorsal position relative to PL1 and depth. PL9 is a landmark that is characterized by its large size, visible through the most posterior sections of the antennal lobe. PL9 is positioned laterally diagonal and opposite to the MD1 glomerulus. PL1-9 are spatially invariant glomeruli. Four residual variant glomeruli are sometimes observed in more posterior sections (PL10-PL13). See Figure S1 (j and k) and Figure S2 (j).
12. **Dorsal group**: The D group consist of 4 invariant glomeruli (D1, D2, D4 and D5) and are the most dorsal group of glomeruli in posterior slices. D1, D2 and the typically observed variant glomerulus D3 are arranged in a triangular manner in the female antennal lobe, with D1 found at the apex and D2 and D3 found ventrally. D3 is sometimes observed laterally to D2. In the male, D1, D2 and D3 form a row along the M-L axis. The second group of glomeruli are constituted by D4 and D5. D5 is considered a landmark glomerulus because of its large size and most dorsal position. D4 is found ventral and more medial relative to D5. Two additional variant glomeruli (D6-D7) were identified in some reconstructions in more posterior slices. See Figure S1 (l) and Figure S2 (i).
13. **Medio-Dorsal group**: The MD group typically comprises 3 glomeruli that flank the medial margin of the antennal lobe. These glomeruli are positioned in the most posterior slices of the antennal lobe from −28 µm to −35 µm, and are named MD1, MD2 and MD3. MD1 is the most posterior and largest glomerulus in the antennal lobe, situated at the same depth as the dorsal neuropil including the ellipsoid and the fan shaped bodies, on either side of the esophagus. The landmark MD1 forms the core of this group. MD2 and MD3 are more anterior but are positioned around MD1 like satellite glomeruli. MD3 is the most anterior and dorsal of these glomeruli. A variant glomerulus MD4 is typically observed anterior to MD1. The MD glomeruli are known to receive input from the maxillary palp neurons (12). An easily identifiable array of posterior antennal lobe glomeruli with increasing depth is: PL9-MD2-V8-MD1, with PL9, MD1 and V8 as landmarks. PL9 and MD1 have a distinct shapes and large surface areas. See Figure S1 (m) and Figure S2 (i and j).

Representative confocal stacks and color-coded 2D maps detailing the typical spatial arrangement of invariant and variant glomeruli in the female left and right antennal lobes as shown by nc82 staining, are illustrated in Figure 2 and Figure S3, respectively. Examples for the left and right antennal lobes of adult males are illustrated in Figure 3 and Figure S4, respectively. The gross 3D shape of reconstructed nc-82 stained antennal lobes varied noticeably between individual mosquitoes (see Figure S5). However, the eight characteristic landmark glomeruli could reliably be identified in all reconstructed models we generated, facilitating assignment of names to all remaining glomeruli based on their relative spatial position.

**Figure 2.**
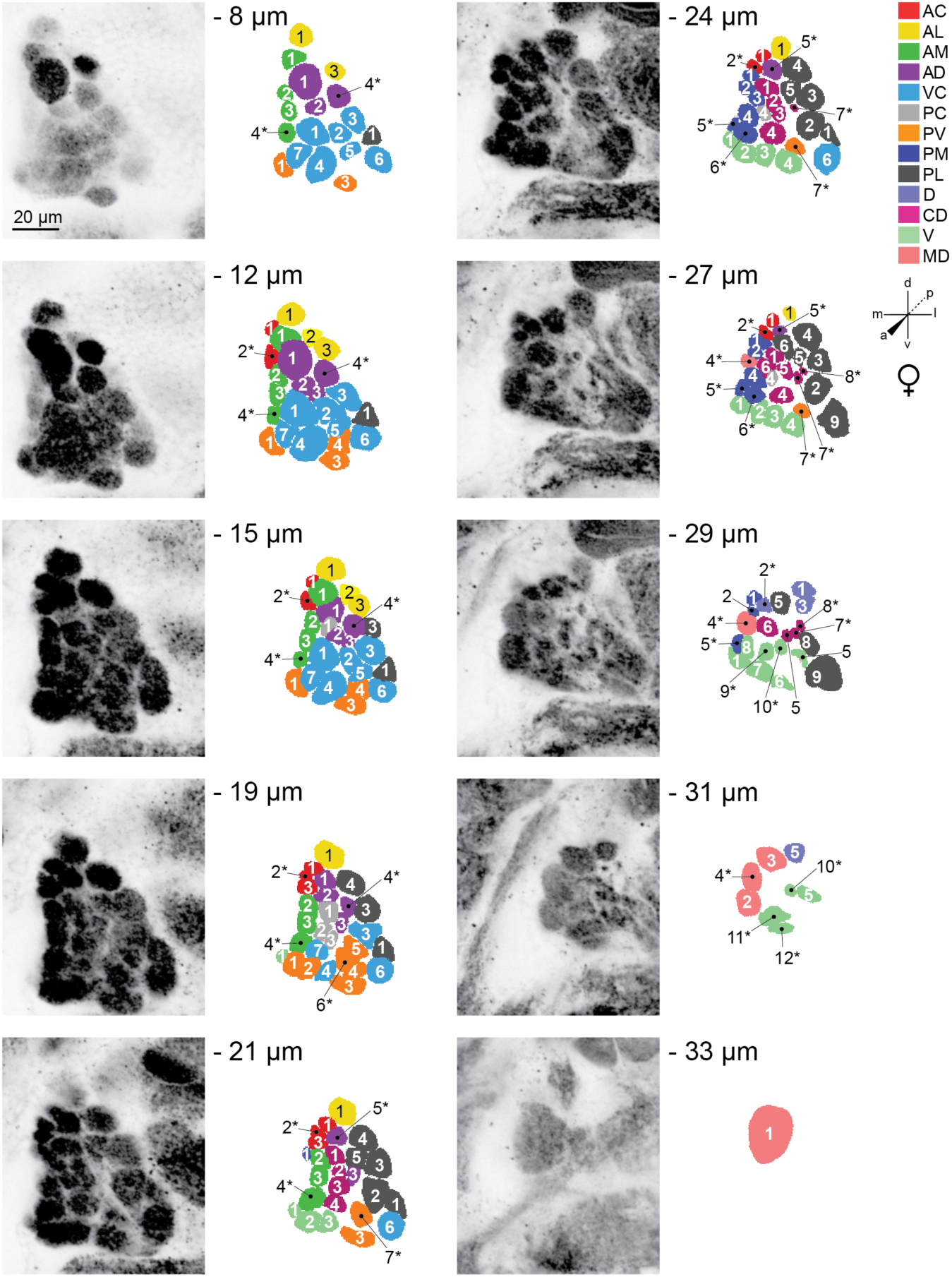
Representative confocal stack from the left antennal lobe of an adult female LVPib12 *Aedes aegypti* as shown by nc82 staining. Ten frontal planes from a total of 35 images taken at 1µm intervals were selected for illustration of the typical geometric arrangement of glomeruli; Scale bar, 20 µm. The depth of each confocal slice is indicated. Glomeruli within each reconstructed slice are color-coded according to their predicted spatial group. Glomeruli are numbered, with 61 out of 63 spatially invariant glomeruli evident in the ten antennal lobe slices depicted here. Variant glomeruli are indicated by asterisks.

**Figure 3.**
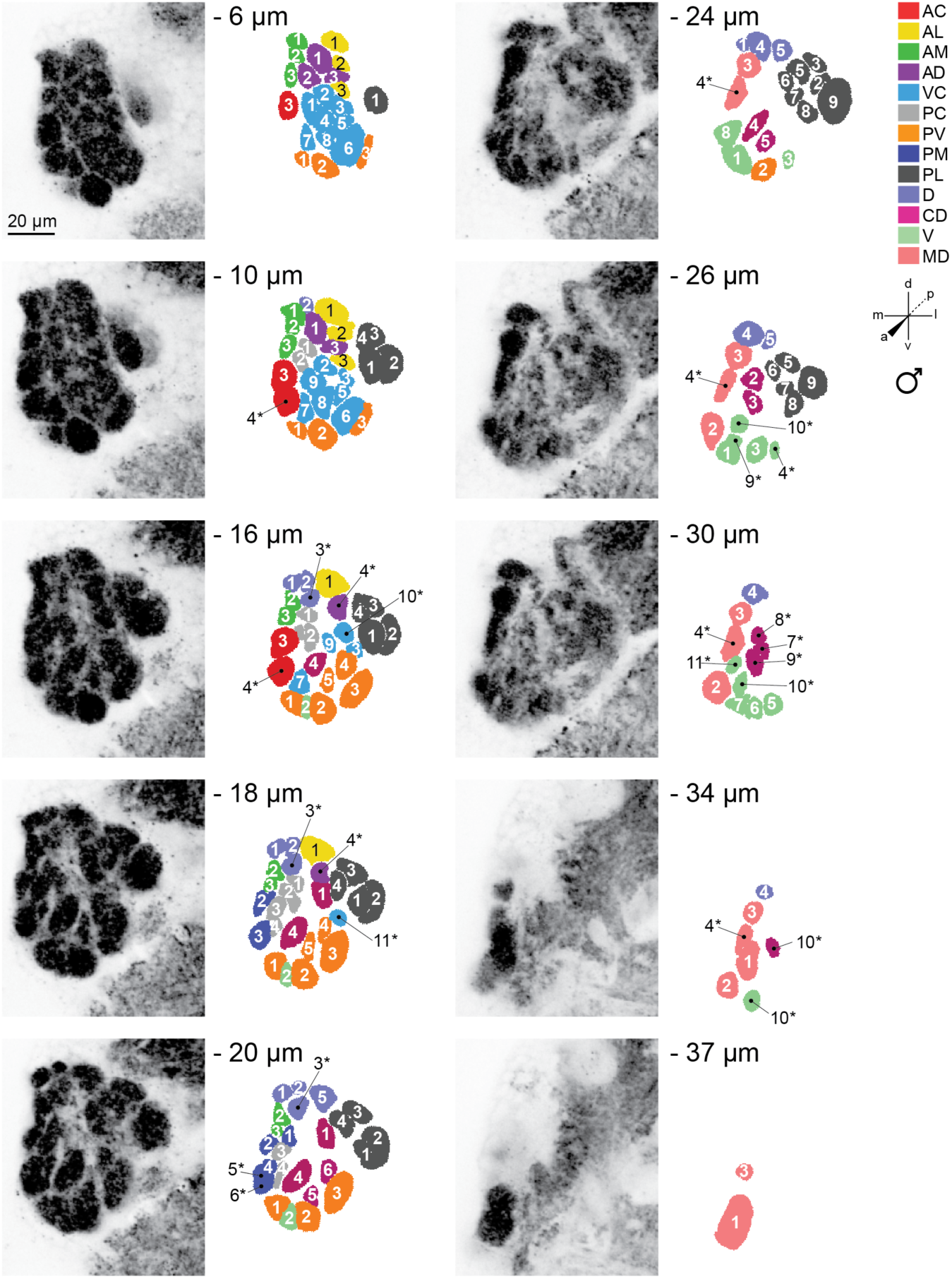
Representative confocal stack from the left antennal lobe of an adult male LVPib12 *Aedes aegypti* as shown by nc82 staining. Ten frontal planes from a total of 40 images taken at 1µm intervals were selected for illustration of the typical geometric arrangement of glomeruli; Scale bar, 20 µm. The depth of each confocal slice is indicated. Glomeruli within each reconstructed slice are color-coded according to their predicted spatial group. Glomeruli are numbered, with 59 out of 63 spatially invariant glomeruli evident in the ten antennal lobe slices depicted here. Variant glomeruli are indicated by asterisks.

### Changes in glomerular nomenclature relative to previous *in vitro* models of the *Aedes aegypti* antennal lobe

We used our reference key (Table 1) to apply our updated system for glomerular nomenclature to the extant *in vitro* models of the *Ae. aegypti* antennal lobe reported by Ignell *et al*. (2005). These authors previously annotated 50 and 49 glomeruli in female and male antennal lobe of the Rockefeller *Ae. aegypti* strain, respectively.

In total, we identified 40 glomeruli in their synapsin:phalloidin-based model of the female antennal lobe (11) and 39 glomeruli in their nc82-based model of the male antennal lobe (11) that appear to occupy orthologous positions to spatially invariant glomeruli annotated in this study. Using our updated system for glomerular nomenclature, only 7 of these glomeruli retain their previously annotated name in both sexes across both studies (AL1-3, AM1-3 and MD1), with the names for all other glomeruli changing (Table S2).

Of those renamed as reported in Table S2, 23 glomeruli in their female model and 24 glomeruli in their male model were reclassified into new glomerular spatial groups with different nomenclature and revised organization. The remaining 10 renamed glomeruli in their female model and 8 renamed glomeruli in their male model were assigned new numerical descriptors to reflect their arrayed position within their extant spatial group – predominantly belonging to the PL, CD and MD groups in both sexes. Across all spatially invariant glomeruli we identified within their models, only 14 (35-36%) share the same descriptor annotated by Ignell *et al*. (2005) in both sexes (Table S2), consistent with considerable variation in the way glomeruli were ascribed names by these authors in their reported models with reference to relative spatial position.

In addition to the complement of orthologous glomeruli described above, 12 new spatially invariant glomeruli in females and 13 new spatially invariant glomeruli in males were identified in our updated atlas that likely result from segmentation of the Johnston’s Organ Center (JOC) from Ignell et al. (2005) (Table S2). We further annotated 11 additional new spatially invariant glomeruli in the updated PL, D, CD, V and MD groups of females, and 11 new spatially invariant glomeruli in the updated PC, PL, CD, V and MD groups of males in our atlas that were not evident in the reconstructed maps of Ignell *et al*. (2005) (Table S2).

According to our reference key, in the female model reported by Ignell *et. al*. (2005), 7 glomeruli are likely spatially variant glomeruli and include those annotated by these authors as V4, AM5, PM2, PM3, MD3, PD3 and PD4 (Table S3). In the male model, 9 glomeruli are likely spatially variant glomeruli and include those annotated by these authors as AL4, AM4, AM7, AC3, AC4, PM1, PM3, PM4 and MD3 (Table S4).

In the antennal lobe models reported by Ignell et al. (2005), we were unable to assign updated names to PC1 and PL6 for the female (Table S3), and additionally AD4 for both sexes (Table S3 and S4). These glomeruli appear positioned at the boundary of two different spatial groups, therefore making definitive assignment to either group difficult. AD4 for example, is positioned at the interface of updated PC and PM groups; PC1 could be assigned either to the updated PC or CD groups, and similarly, PL6 could be considered as an updated V or PL group glomerulus.

### Volumetric analysis of the *Aedes aegypti* antennal lobe

With our updated system for glomerular nomenclature established, we then performed volumetric analyses on the left and right antennal lobes from all nc82-stained LVPib12 brains that we reconstructed (*n* = 5 brains per sex, 20 antennal lobes total in this dataset). The mean total antennal lobe volume is 123,319 µm^3^ in females, and 85,370 µm^3^ in males (*Mann-Whitney*, *P*<0.0001). The total antennal lobe volume of females is therefore approximately 1.4 times larger than that of male mosquitoes.

For both sexes, there were no significant differences in total antennal lobe volume between the left and right sides of the brain (Table S5). Within antennal lobes derived from females (Figure 4) and males (Figure 5), the mean volumes of glomeruli varied significantly (one-way ANOVA, *P*<0.0001 for each lobe – Figures S6-S9). The largest glomerulus in both the female and male antennal lobe is MD1, with a mean volume of 5,222 µm^3^ and 4,090 µm^3^, which accounts for 4.26% and 4.81% of their total antennal lobe volumes, respectively.

**Figure 4.**
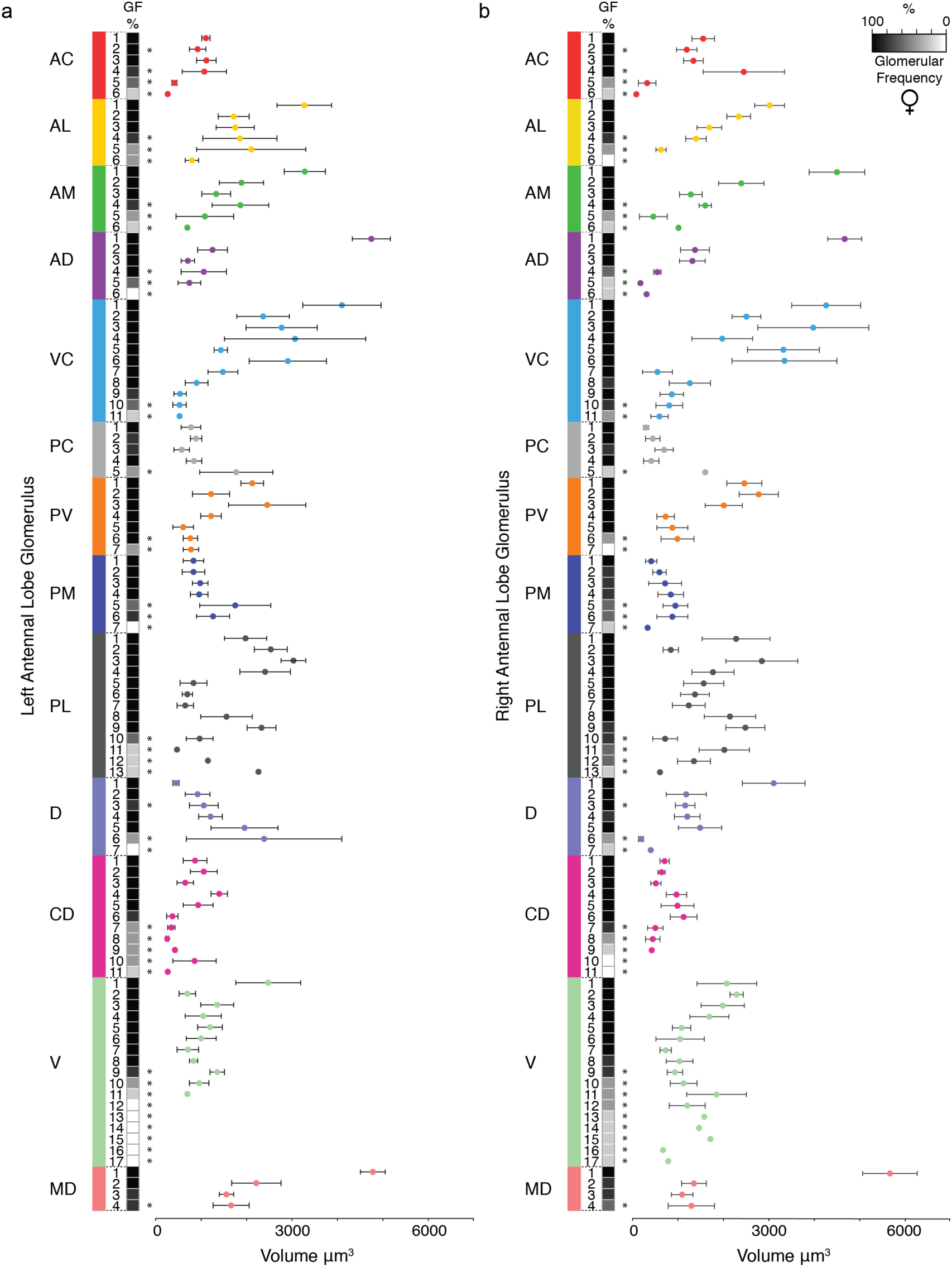
Frequency and volume of glomeruli observed in three-dimensional reconstructions of the (**a**) left and (**b**) right antennal lobe of adult female LVPib12 *Aedes aegypti* with nc82 staining. A heatmap of glomerular frequency (GF%) and the associated classification of glomeruli assigned to each of the 13 glomerular spatial groups during replicate reconstructions is shown. Invariant glomeruli are classified as those having a GF of ≥80% across all antennal lobe reconstructions, while variant glomeruli below this threshold are indicated by asterisks (*n* = 5 brains).

**Figure 5.**
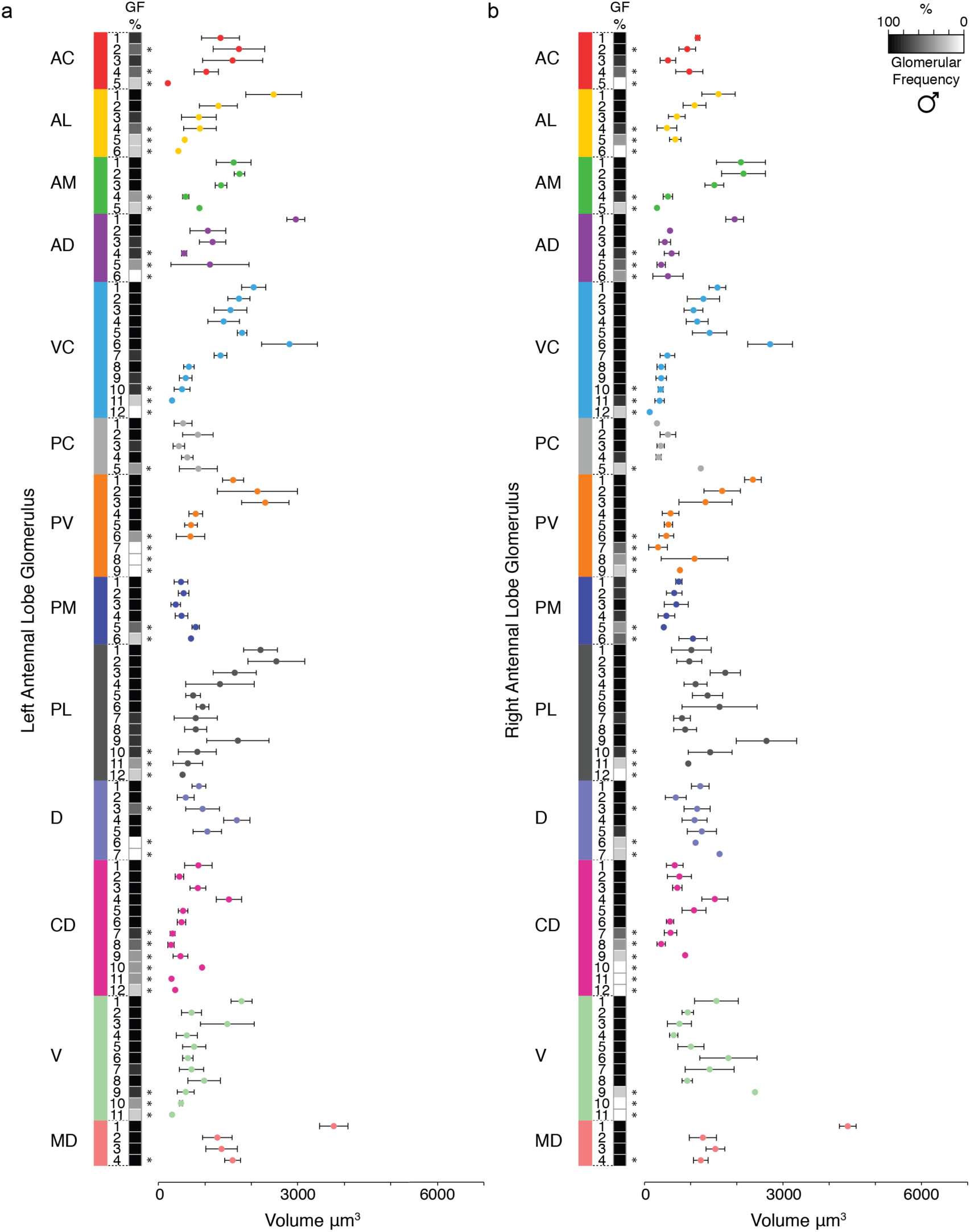
Frequency and volume of glomeruli observed in three-dimensional reconstructions of the (**a**) left and (**b**) right antennal lobe of adult male LVPib12 *Aedes aegypti* with nc82 staining. A heatmap of glomerular frequency (GF%) and the associated classification of glomeruli assigned to each of the 13 glomerular spatial groups during replicate reconstructions is shown. Invariant glomeruli are classified as those having a GF of ≥80% across all antennal lobe reconstructions, while variant glomeruli below this threshold are indicated by asterisks (*n* = 5 brains).

To identify candidate sexually dimorphic glomeruli, we then calculated relative volumes for all 63 invariant glomeruli that we annotated in females and males (Table S5). To do this, we determined the percentage volume that each invariant glomerulus sampled occupies within its concordant antennal lobe, and then compared grouped relative glomerular volumes by sex using the non-parametric *Mann-Whitney U*-Test. Using this approach, we identified two glomeruli with significantly larger relative volumes in females: AD1 (*P* = 0.005) and VC1 (*P* = 0.036). Of these, the landmark glomerulus AD1 has a mean volume of 4,708 µm^3^ in females and 2,456 µm^3^ in males which accounts for 3.82% and 2.89% of their total antennal lobe volumes, respectively. In addition, four glomeruli were significantly larger in males: AM3 (*P* = 0.004), CD3 (*P* = 0.019), CD4 (*P* = 0.015) and MD3 (*P* = 0.016). AM3, which is positioned medially on the anterior surface of the antennal lobe, has a mean volume of 1,315 µm^3^ in females and 1,431 µm^3^ in males which accounts for 1.06% and 1.67% of their total antennal lobe volumes, respectively.

### Validating the revised neuroanatomical model of the *Aedes aegypti* antennal lobe

To verify glomerular counts and the spatial organization of the antennal lobe model reconstructed from nc82-staining, we utilized a complementary and independent neuronal labeling method to visualize glomeruli. Since the antennal lobes across diverse insect species are known to contain a rich aggregation of F-actin filaments in OSNs (23), the fluorophore-conjugated toxin phalloidin was used as a cytoskeletal marker of OSN axonal processes to visualize antennal lobe morphology. Using this method, we could observe strong staining of all the major neuropil in the central *Ae. aegypti* brain (Figure 6a and 6b). We then performed reconstructions on the left and right antennal lobe from 3 male and 3 female LVPib12 mosquitoes. We counted 79 ± 1 glomeruli in females and 77 ± 3 glomeruli in males (mean ± s.d.) (Figure 6g and 6h).

**Figure 6.**
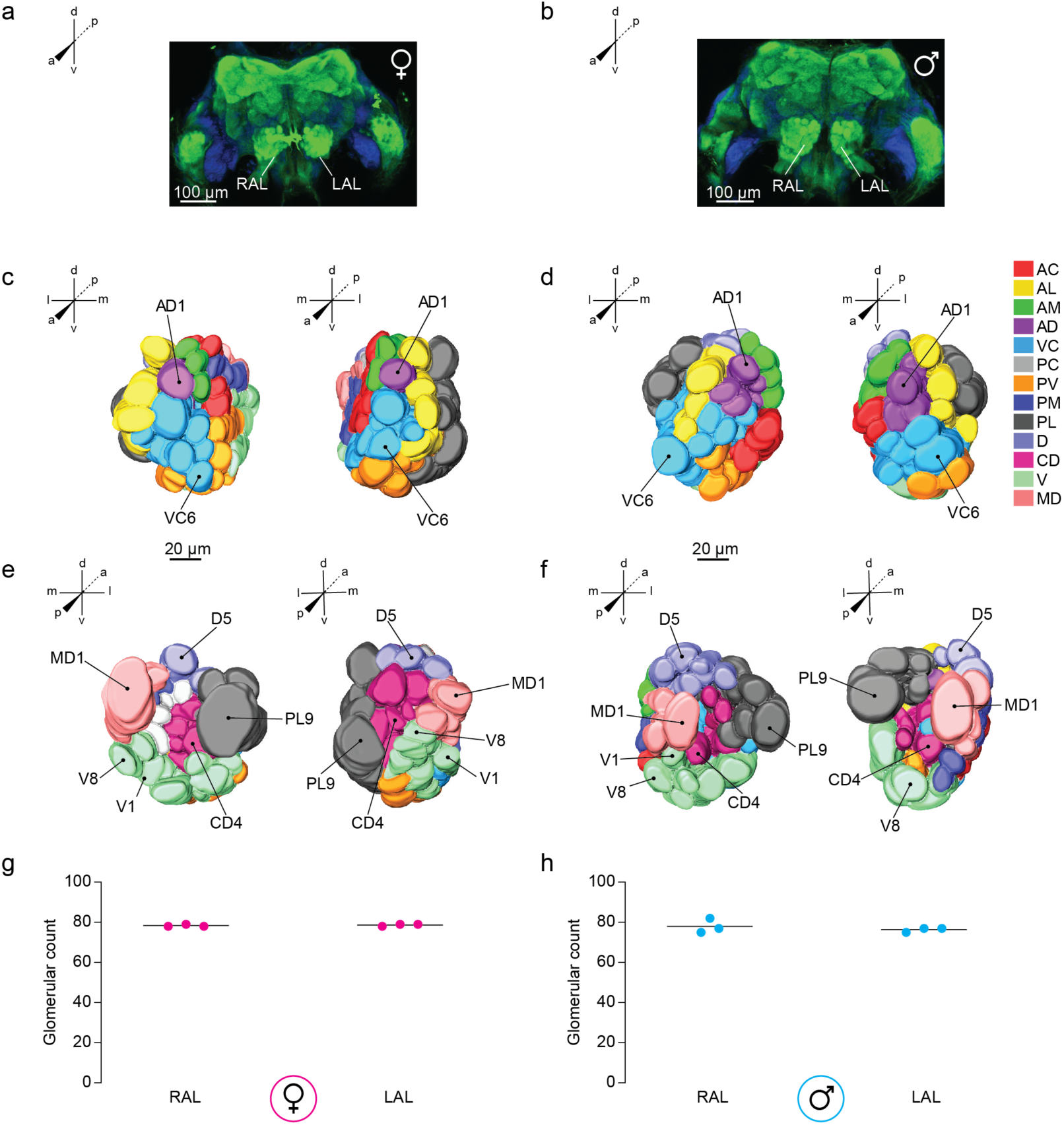
Antennal lobe morphology of the LVPib12 *Aedes aegypti* strain as shown using phalloidin staining. Confocal images of a representative adult (**a**) female and (**b**) male brain with F-actin filaments labeled with a phalloidin-conjugated fluorophore (green) and nuclei labeled with DAPI dye counterstain (blue); Scale bars, 100 µm. Right (RAL) and left antennal lobes (LAL) are indicated. Anterior and posterior views of three-dimensional reconstructive models of the right and left antennal lobe from a female (**c** & **e**) and male (**d** & **f**) mosquito are shown, respectively. Visible landmark glomeruli on these surfaces are denoted. Antennal lobe glomeruli are color-coded according to their classification into 13 groups based on their spatial position along anterior-posterior (a-p), dorsal-ventral (d-v), and medial-lateral (m-l) co-ordinate axes. The total number of glomeruli found in reconstructed (**g**) female and (**h**) male antennal lobes are shown (*n* = 3 brains per sex).

Relative to pre-synaptic nc82-staining, the diffuse labeling pattern of phalloidin from overlapping OSN axonal processes traversing the antennal lobe made delineation of glomeruli using this method challenging at times, particularly in the anterior and central antennal lobe. As a result, the confocal images had to be magnified 4-5 times in order to perform segmentation. Furthermore, at lower magnification we observed in many reconstructions that PV and VC glomeruli appeared to be stained as a single large cluster of neuropil, rather than as individual glomerular units that could be clearly demarcated with antibody labeling. This cluster has a similar morphology to the non-glomerular JOC that was described in an earlier *Ae. aegypti* antennal lobe model generated using phalloidin staining (11).

We selected representative reconstructions of the male and female antennal lobes in which we were able to easily demarcate all landmark glomeruli, as described for the nc82-stained model (Figure 6c to 6f). We then created 2D anatomical maps of female left and right antennal lobes (Figure S10 and S11) to comprehensively name all glomeruli in this dataset. Consistent with the nc82 model, we found that the antennal lobe was divisible into 13 distinct spatial groups following our reference key. Applying our thresholding criteria to categorize spatially invariant and variant glomeruli within this data set, we identified the exact complement of spatially invariant glomeruli (n=63) seen with nc82-stained brains. Remaining glomeruli found at a frequency of <80% in the antennal lobe of both sexes were categorized as variant glomeruli, consistent with morphological distortions being inherent to both nc82 and phalloidin-based neuronal staining methods. Of note, the PL group of glomeruli was positioned more dorsally in reconstructions of the male left antennal lobe (Figure 6f).

To further assist with user identification of glomeruli in the LVPib12 strain using nc82 and phalloidin-based neuronal staining methods, we have generated a series of digital models that allow users to interactively view the spatial arrangement of glomeruli in left antennal lobe of a female brain sample stained with each method from different perspectives and depths (Supplementary Movie S1-S4).

### Conservation of antennal lobe organization across geographically divergent strains of ***Aedes aegypti***

Finally, we compared the antennal lobe neuroanatomy of the inbred LVPib12 strain (21) originally of African origin, with a recently established strain of *Ae. aegypti* derived from Patillas, Puerto Rico (Figure 7a) that is routinely supplemented with field-caught mosquitoes (24). We applied nc82-staining to dissected and fixed Patillas brain samples and reconstructed the left and right antennal lobes of three male and three female *Ae. aegypti* mosquitoes in total. Overall, the gross shape of antennal lobes from the LVPib12 and Patillas strains differed subtly. In particular in the Patillas strain, the female left and right antennal lobes were rounded, while the PV group from the male left and right antennal lobes appeared to taper out of the central region. Using our LVPib12 antennal lobe models as a guide, we generated 3D models of the Patillas male and female antennal lobe, in which we classified and color-coded glomeruli into the 13 glomerular spatial groups (Figure 7b to 7e).

**Figure 7.**
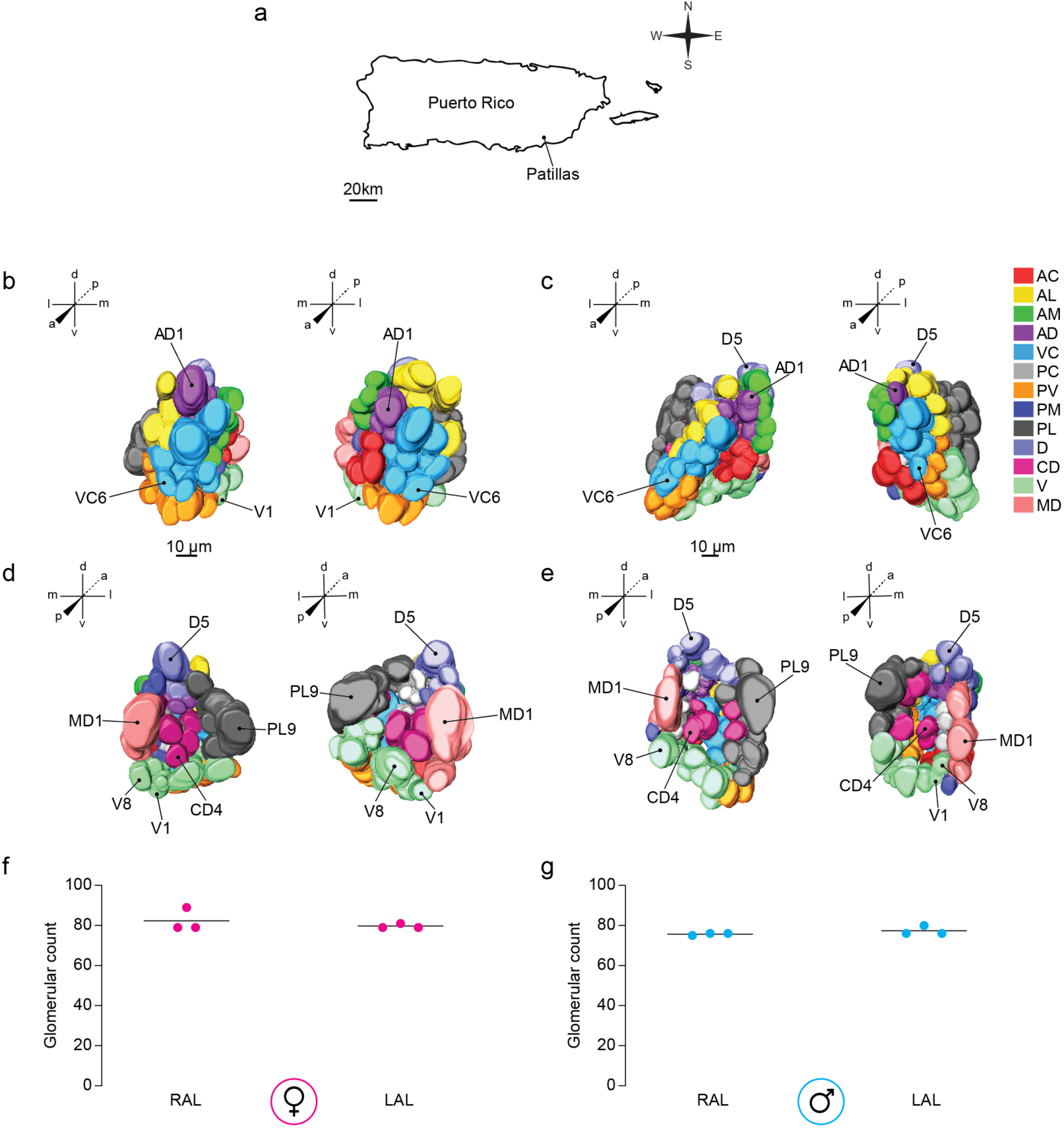
Antennal lobe morphology of the Patillas *Aedes aegypti* strain as shown using nc82 staining. This mosquito strain was founded with field-collected *Ae. aegypti* from the municipality of Patillas, Puerto Rico (**a**). Anterior and posterior views of three-dimensional reconstructive models of the right and left antennal lobe from a female (**b** & **d**) and male (**c** & **e**) mosquito are shown, respectively. Visible landmark glomeruli on these surfaces are denoted. Antennal lobe glomeruli are color-coded according to their classification into 13 groups based on their spatial position along anterior-posterior (a-p), dorsal-ventral (d-v), and medial-lateral (m-l) co-ordinate axes. The total number of glomeruli found in reconstructed (**f**) female and (**g**) male antennal lobes are shown (*n* = 3 brains per sex).

We then segmented 3D models from a representative Patillas male and female brain sample to yield 2D maps to name individual glomeruli. Using our reference key (Table 1), we named all antennal lobe glomeruli in these brain samples, and then applied our thresholding criteria to identify spatially invariant and variant glomeruli within the data set. We identified all 63 spatially invariant glomeruli found in the LVPib12 strain and denoted the presence of residual variant glomeruli consistent with our previous models. In total, we counted 81 ± 4 glomeruli in the female antennal lobes (Figure 7f) and 77 ± 2 in the male antennal lobes (mean ± s.d.) (Figure 7g). We conclude that despite gross morphological differences in antennal lobe shape, the spatial organization of antennal lobe glomeruli in these geographically diverse populations of *Ae. aegypti* appears conserved.

## Discussion

*Aedes aegypti* is an emerging model system for mosquito olfaction (3, 25–31). An accurate anatomical map of the antennal lobe is crucial towards studies aimed at understanding how odors are detected and represented by the olfactory system of this disease vector. In this study, we generated an updated atlas for the *Ae. aegypti* antennal lobe providing systematic nomenclature and a standardized framework for identifying glomeruli within this primary olfactory processing center of the mosquito brain.

We determined the *Ae. aegypti* antennal lobe is composed of approximately 80 morphologically discrete glomeruli in both sexes. We surmise that the advanced confocal imaging techniques employed in our study may have facilitated clearer delineation of glomerular boundaries, thus revealing additional glomeruli relative to previous studies that found 35 or 49-50 glomeruli in the *Ae. aegypti* antennal lobe, respectively (11, 18). In comparison with the *Drosophila melanogaster* antennal lobe which has 56 glomeruli (32), the *Aedes aegypti* antennal lobe has markedly increased subdivision making assignment of names to individual glomeruli not a trivial matter.

To facilitate stereotypical identification of glomeruli, we first subdivided the antennal lobe into 13 spatial groups with revised nomenclature, merging several extant groups from a previous antennal lobe model (11), and annotating others with new descriptors to better reflect their true spatial position and accommodate the expanded complement of glomeruli we found (Table S1). Subsequently, we identified eight landmark glomeruli based on their characteristic shape, volume and position across these groups, with all other glomeruli annotated by their spatial position relative to these landmarks. Guided by replicate morphological reconstructions, we annotated 63 spatially invariant glomeruli that can be reliably identified in the same geometric arrangement within dissected and stained antennal lobe preparations.

Despite intra- and inter-strain variability in the overall shape of antennal lobes in this study, the spatial organization of invariant glomeruli from LVPib12 and Patillas strains was shared and discernable. These results suggest that antennal lobe organization is likely conserved across geographically divergent strains of *Ae. aegypti*. Our revised atlas may therefore serve as a useful reference for studying antennal lobe organization in other *Ae. aegypti* strains supplemental to those reported here.

We determined the largest invariant glomerulus in *Ae. aegypti* was MD1. Anterograde tracing experiments in this species previously revealed that MD1 likely receives projections from maxillary palp-derived OSNs (12) which are responsive to the volatile gas carbon dioxide (CO_2_). The increased volume of this glomerulus in both sexes may be reflective of the critical role that CO_2_ plays in *Ae. aegypti* host-seeking behavior (3, 30, 33). We additionally identified six candidate glomeruli including AD1 and AM1 with sex-specific differences in relative volume. In the future, identifying what odorants these glomeruli detect may help to further clarify their role in sexually dimorphic olfactory behaviors in this mosquito species.

We noted considerable variation in the spatial position of approximately 20% of glomeruli annotated within our *in vitro* antennal lobe reconstructions. These variant glomeruli tended to have smaller volumes and were clustered around boundaries between spatial groups, making them difficult to accurately name. Typically, variant glomeruli were positioned in more posterior layers of the antennal lobe and were frequently noted in PM, PL, CD and V groups. Such spatial variability in glomerular position is likely due to technical artifacts associated with dissection, fixation, protocol duration and coverslip placement for imaging that commonly distort antennal lobe morphology (34). The *Ae. aegypti* antennal lobes are also positioned in close proximity to the anterior esophagus, a thick structure lined with cuticle, epithelial cells and muscle (35). The esophagus runs through the esophageal foramen and it was removed during our dissections to prevent it from occluding the antennal lobes, possibly causing distortion or disassociating glomeruli, and altering the shape of each antennal lobe.

As recently described for a revised antennal lobe model for *Anopheles coluzzii* (20), we did not observe any large clusters of neuropil orthologous to the Johnston’s Organ Center (JOC) (11) in any of our reconstructions from the LVPib12 or Patillas *Ae. aegypti* strains. Instead, we identified morphologically discrete glomeruli in the anterior region of the antennal lobe where this structure was previously annotated. The JOC was assigned based on evidence from silver nitrate Golgi staining experiments (15) that suggested putative neuronal connectivity between the antennal lobe and the sound-receptive Johnston’s Organ (JO). However, other anterograde tracing studies in *Ae. aegypti* and *D. melanogaster* indicate afferent JO neurons project to the antennal motor and mechanosensory center (AMMC) of the deutocerebrum, rather than the antennal lobe (36, 37). Further in-depth neuroanatomical studies are clearly warranted to definitively validate any afferent or efferent neuronal connectivity between the JO and antennal lobe in *Ae. aegypti* and other mosquito species.

The *Ae. aegypti* genome encodes large chemoreceptor gene families including Odorant Receptors, Ionotropic Receptors and Gustatory Receptors (28, 38). To date, chemoreceptors have only been localized to OSNs on the *Ae. aegypti* antennae, maxillary palps and proboscis using bulk transcriptomics and immunohistochemical approaches (25, 28, 39). We propose this model atlas will be useful to map projection patterns of neurons expressing chemoreceptors and other genes involved in olfactory signaling to their associated glomeruli in the *Ae. aegypti* antennal lobe using genetic labeling approaches (20, 34, 40, 41). Such studies will be critical to validate our antennal lobe model that is currently based on morphology alone, toward understanding the molecular and cellular organization of the *Ae. aegypti* olfactory system.

Finally, this updated atlas lays the foundation for improved spatial registration during functional imaging within the *Ae. aegypti* antennal lobe with genetically encoded calcium indicators and dyes (30, 42, 43). In particular, studies with activity-dependent ratiometric indicators such as CaMPARI (44, 45), that provide a permanent readout of neural activity and are also compatible with post-hoc neuronal staining to aid in spatial registration of such signals, will benefit greatly from using the *in vitro* antennal lobe atlas reported here as a reference. Such analyses may help to reliably identify glomeruli activated or inhibited by diverse odorants that are behaviorally relevant to *Ae. aegypti* such as human odors, plant odors, pheromones and insect repellents. This information may further accelerate reverse engineering of synthetic odorant blends mimicking patterns of neurophysiological activity elicited by these natural olfactory stimuli, to yield novel applied strategies to modulate *Ae. aegypti* olfactory behaviors.

## Materials and Methods

### Mosquito maintenance and rearing

*Aedes aegypti* LVPib12 (21) and Patillas (24) strains were maintained at 27°C, 80% relative humidity with a 12 hr light:dark photoperiod. Immature stages were reared using a standardized protocol optimized for yielding synchronous pupation. Briefly, on day 0 (hatching), 500mL of a 1% w/v autoclaved broth solution of finely milled TetraMin (Tetra, 16110) in dH_2_O was placed in a clean polycarbonate pan (Cambro, 24CW135) with covering lid (Cambro, 20CWCH135). Oviposition papers containing desiccated eggs were then completely submerged in the broth solution inside each allocated pan to facilitate larval hatching. On day 1, 500 ml of additional dH_2_O and 1g of finely milled Tetramin was added to the pan to bring the total volume to 1L. On day 2, oviposition papers were removed, and hatched larvae thinned to a set density of 200 larvae in 2L of fresh dH_2_O per pan and fed one coarsely dissociated 1g Tetramin tablet. On day 3, larvae in each pan were fed with an additional 1g Tetramin tablet. Thereafter, larvae in each pan were fed two 1g Tetramin tablets per day until pupation which fully occurred on day 6. After emergence, adult mosquitoes were provided constant access to a 10% w/v sucrose solution. For routine colony maintenance, females were blood fed using anesthetized mice (Johns Hopkins Animal Care and Use Committee protocol # MO18H391).

### Neuronal staining

Neuropil in the central brain of mated, 5-10 day old, male and nulliparous female *Ae. aegypti* were initially stained with the primary monoclonal antibody nc82 (DSHB, nc82-s, AB_2314866) that labels the pre-synaptic active zone protein Bruchpilot. To dissect brains, mosquito heads were severed from the thorax and tissue fixed for 3 hr at 4°C in Milonig’s buffer: 0.1M PBS pH 7.2 with 4% v/v paraformaldehyde (PFA) as a fixative. Next, adult brains were dissected out into 0.1 M PBS using fine forceps (Dumont No 5, 100 nm tips) to carefully remove the head capsule as well as the pigmented ommatidia over the optic lobes and any floating air sacs connected to the brain. Dissected brains were washed 3 times for 20 min each at room temperature in 0.25% PBST consisting of 0.25% v/v Triton-X-100 detergent as a permeabilizing agent prepared in 0.1 M PBS. The brains were then treated in a blocking solution consisting of 2% w/v normal goat serum (NGS) and 4% v/v PFA in 0.1 M PBS and incubated on a nutator overnight at 4°C. This step was followed by incubation in a primary antibody solution for 3 days at 4°C consisting of mouse anti-nc82 (1:50 v/v in 0.1 M PBS with 0.25% Triton-X and 2% NGS). Next, the brains were washed three times at room temperature in 0.25% PBST and incubated for 3 days at 4°C in a secondary antibody solution consisting of goat anti-mouse Cy3 (Jackson ImmunoResearch 115-165-062, 1:200 v/v in 0.1 M PBS with 0.25% Triton-X and 2% NGS). Following three final washes at room temperature for 20 min each in 0.25% PBST, the brains were mounted within the space created by two coverslip (Number 2 - 170 µm) bridges on glass slides in 20µl of Slow-Fade Gold Antifade Mountant (Invitrogen, S36936), to preserve 3D structure.

As an alternative method to immunohistochemistry, adult brains were stained with fluorophore conjugated-toxin phalloidin to mark cytoskeletal F-actin filaments present ubiquitously across neuropilar structures including OSN axonal processes. Adult heads were first fixed in Milonig’s buffer for 3 hr at 4° C and brain tissue dissected in 0.1 M PBS as described above. After washing 3 times in 0.25% PBST, the dissected brains were incubated in 0.25% PBST with 1/80 (v/v) Alexa Fluor 488 Phalloidin (Invitrogen, A12379) and 2µg/ml DAPI nuclear counterstain (Invitrogen, D1306), for 3 days at 4° C. Brains were then mounted on glass slides as described and imaged.

### Image acquisition settings

Images of nc82- and phalloidin stained brains were taken on a single-point laser scanning, Carl-Zeiss LSM 780 confocal microscope. To capture images of the entire adult brain for both methods, a 10X objective lens (0.3 NA, Plan-Apochromat) was used. To yield images to perform 3D reconstructions of the antennal lobes, images were taken with a 20X objective lens (0.8 NA, Plan-Apochromat). Images were processed using Imaris software (Oxford Instruments).

For nc82-stained brains, a solid-state laser line at 561 nm (1% laser power) was used to excite the Cy3 signal and the gain of the GaAsP detector was set to 600. 100 z-slices, with a z-step size of 1 μm and a 1024 X 1024 pixel size were acquired. For 3D reconstructions, 60 z-slices with a z step size of 1 μm were acquired. The power of the 561 nm laser was adjusted to 2% with detector gain at 620. These settings ensured the acquisition of high intensity images, allowing for the accurate de-lineation of glomerular boundaries while scanning through the entire volumes of both antennal lobes, including a few slices above the surface of the antennal lobe, and a few slices below the deep-seated MD1 glomerulus, so as to closely replicate three-dimensionality of the AL structure in the reconstructed model. Images were imported in the *.lsm-Zeiss, Zen format into the Amira software program. We generated inverted images of the z-slices, in Image j, first using the gray lookup table (LUT) to get a black and white image, followed by the invert LUT. No brightness and contrast adjustments were made.

For phalloidin stained brains, a 488 nm laser line (2% laser power) was used to excite the Alexa-488 phalloidin fluorophore, with the detector gain at 500. The DAPI stain was excited with a 405 nm diode laser (2% laser power and detector gain at 400). 60 z-slices with a z step size of 1 μm and a 1024 X 1024 pixel size were acquired. Phalloidin and DAPI stained confocal slices were inverted using Adobe Photoshop, without altering brightness or contrast.

### Antennal lobe reconstructions

3D reconstructions of the antennal lobe were performed using Amira software (FEI Houston Inc). Confocal files were opened in the ‘.lsm’ format and viewed using the Image Segmentation Editor tool. For each z-slice typically ranging from 0 µm to −60 µm through the anterior-posterior axis of the antennal lobe, olfactory glomeruli were identified by shape and position. We defined the start of the antennal lobe (denoted 0µm) as the first z-slice where the surface of the first landmark and most anterior glomerulus AD1 is located. Next, an image segmentation step was carried out in which the voxels occupied by individual glomeruli were first highlighted using the Amira paintbrush tool and then assigned a unique label and color. After segmentation of the voxels corresponding to all antennal lobe glomeruli, the image volume was surface rendered to create a 3D model to allow visualization of antennal lobe morphology along x-y-z axes.

Volumes of individual glomeruli were obtained (in µm^3^) using the Material Statistics tool in Amira. We used the ortho slice tool to generate 2D cartoons of all the confocal slices and saved the images in ‘tif’ format. Each antennal lobe glomerulus was named based on its spatial position relative to other glomeruli. For each strain and staining method used, we classified glomeruli as spatially ‘invariant’ or ‘variant’ based on their frequency of identification across combined male and female, left and right antennal lobe datasets. A threshold frequency of ≥80% was designated for the classification of spatially invariant glomeruli.

### Statistics

Based on our 3D atlas of the antennal lobe of female and male LVPib12 *Aedes aegypti*, we calculated the volume of each annotated glomerulus and the total volume of the antennal lobe (constituting the cumulative volume of all stained glomeruli in a given antennal lobe) from each brain sample. To test whether mean volumes of invariant glomeruli differed significantly within each lobe, we performed a *One-Way ANOVA* and *Tukey’s HSD* post hoc test to correct for multiple comparisons. To evaluate size differences of homologous invariant glomeruli between the left and right antennal lobe and between sexes, the non-parametric *Mann-Whitney U*-test was performed. Statistical analysis was performed using GraphPad Prism Software version 8.1.1 (GraphPad Software, Inc.). *P*< 0.05 were considered significant.

## Acknowledgements

We thank Christopher Potter for initial access to Amira software; Roberto Barrera (CDC Dengue Branch, San Juan, PR) for the Patillas stain; Darya Task and Benjamin Burgunder for expert assistance with brain immunohistochemistry and advice on image reconstructions; and Olena Riabinina and members of the McMeniman and Potter laboratories for helpful discussions and comments on the manuscript. This research was supported by funding from the National Institutes of Health NIAID (R21 AI139358-01), Centers for Disease Control and Prevention (200-2017-93143) and USAID (AID-OAA-F-16-00061) to C.J.M. Microscopy infrastructure at Johns Hopkins School of Medicine Microscope Core Facility used in this research was supported by the National Institutes of Health NCRR (S10OD016374). We further acknowledge generous support to the McMeniman laboratory from the Johns Hopkins Malaria Research Institute and Bloomberg Philanthropies. We thank the Developmental Studies Hybridoma Bank at University of Iowa for supplying the monoclonal nc82 antibody generously donated by Erich Buchner.

## Data Availability

Raw imaging files and datasets analyzed in this study are available from the corresponding author upon request.

---

**Table S1:** Changes to nomenclature and/or organization of glomerular spatial group classifiers in this updated version of the *Aedes aegypti* antennal lobe atlas

| Spatial Group – Updated Atlas<sup>1</sup> | Spatial Group – Ignell *et al.* (2005) | Justification for Changes |
| --- | --- | --- |
| Antero-Dorsal (AD) | = Ventral (V) | Renamed to more accurately reflect the anterior and dorsal position of these glomeruli |
| Ventro-Central (VC) | = replaces the JOC | Ventro-Central (VC) and Postero-Ventral (PV) groups replace the previously annotated Johnston's Organ Center (JOC) |
| Antero-Lateral (AL) | = Same name and spatial position | - |
| Antero-Medial (AM) | = Same name and spatial position | - |
| Antero-Central (AC) | = Antero-Medial (AM) + Postero-Medial (PM) | Some glomeruli from the Antero-Medial (AM) and Postero-Medial (PM) groups were found in close proximity in anterior and central sections of the antennal lobe and were reclassified as the updated AC group |
| Postero-Central (PC) | = Antero-Central (AC) + Medio-Central (MC) | The Medio-Central (MC) group and some glomeruli from the Antero-Central (AC) group were consolidated due to their close spatial proximity in posterior and central antennal lobe sections and were reclassified as the updated PC group |
| Postero-Medial (PM) | = Antero-Dorsal (AD) + Antero-Central (AC) + Postero-Dorsal (PD) | Some glomeruli from the Antero-Dorsal (AD), Antero-Central (AC) and Postero-Dorsal (PD) groups were consolidated due to their close spatial proximity in posterior and medial sections of the antennal lobe and were reclassified as the updated PM group |
| Postero-Ventral (PV) | = replaces the JOC | Postero-Ventral (PV) and Ventro-Central (VC) groups replace the previously annotated Johnston's Organ Center (JOC) |
| Ventral (V) | = Postero-Dorsal (PD) | Renamed to more accurately reflect the ventral position of these glomeruli in posterior antennal lobe sections |
| Centro-Dorsal (CD) | = Same name and spatial position | - |
| Postero-Lateral (PL) | = Postero-Lateral (PL) + Latero-Central (LC) | Postero-Lateral (PL) and Latero-Central (LC) groups were consolidated due to their close spatial proximity in posterior and lateral antennal lobe sections |
| Dorsal (D) | = Antero-Dorsal (AD) | Renamed to more accurately reflect the dorsal position of these glomeruli in posterior antennal lobe sections |
| Medio-Dorsal (MD) | = Same name and spatial position | - |

<sup>1</sup> All 13 spatial groups of glomeruli have revised nomenclature and/or organization relative to Ignell *et al.* (2005) as detailed above and in Table S2-S4.

**Table S2.** Summary of nomenclature changes for spatially invariant glomeruli

| Glomerulus<br>name in this atlas | Glomerulus name in<br>Ignell *et al.* (2005) |  |
| --- | --- | --- |
|  | Female | Male |
| AC1 | AM4 | AC2 |
| AC3 | PM1 | AM5 |
| AL1 | AL1 | AL1 |
| AL2 | AL2 | AL2 |
| AL3 | AL3 | AL3 |
| AM1 | AM1 | AM1 |
| AM2 | AM2 | AM2 |
| AM3 | AM3 | AM3 |
| AD1 | V1 | V2 |
| AD2 | V2 | V3 |
| AD3 | V3 | V1 |
| VC1 | JOC | JOC |
| VC2 | JOC | JOC |
| VC3 | JOC | JOC |
| VC4 | JOC | JOC |
| VC5 | JOC | JOC |
| VC6 | JOC | JOC |
| VC7 | JOC | JOC |
| VC8 | JOC | JOC |
| VC9 | JOC | JOC |
| PC1 | AC1 | AC1 |
| PC2 | MC1 | - |
| PC3 | MC2 | - |
| PC4 | MC3 | PC1 |
| PV1 | PM4 | PM2 |
| PV2 | JOC | JOC |
| PV3 | JOC | JOC |
| PV4 | JOC | JOC |
| PV5 | PD8 | JOC |
| PM1 | AD5 | AD5 |
| PM2 | AC2 | AD6 |
| PM3 | PD1 | AD7 |
| PM4 | PD2 | PD1 |
| PL1 | PL5 | PL2 |
| PL2 | PL4 | PL1 |
| PL3 | PL1 | LC1 |
| PL4 | PL2 | - |
| PL5 | LC1 | - |
| PL6 | LC2 | - |
| PL7 | - | - |
| PL8 | PL3 | - |
| PL9 | - | - |
| D1 | - | AM6 |
| D2 | AD3 | AD3 |
| D4 | AD1 | AD1 |
| D5 | AD2 | AD2 |
| CD1 | CD3 | CD1 |
| CD2 | CD4 | CD3 |
| CD3 | CD2 | CD2 |
| CD4 | CD1 | PC2 |
| CD5 | - | PC3 |
| CD6 | - | - |
| V1 | PD5 | PD2 |
| V2 | PD6 | PD3 |
| V3 | PD7 | PD5 |
| V4 | - | PD4 |
| V5 | - | - |
| V6 | - | PD6 |
| V7 | - | PD7 |
| V8 | - | PD8 |
| MD1 | MD1 | MD1 |
| MD2 | - | - |
| MD3 | MD2 | MD2 |

(-) Orthologous spatially invariant glomerulus not identified within model of Ignell *et al.* 2005

**Table S3.**
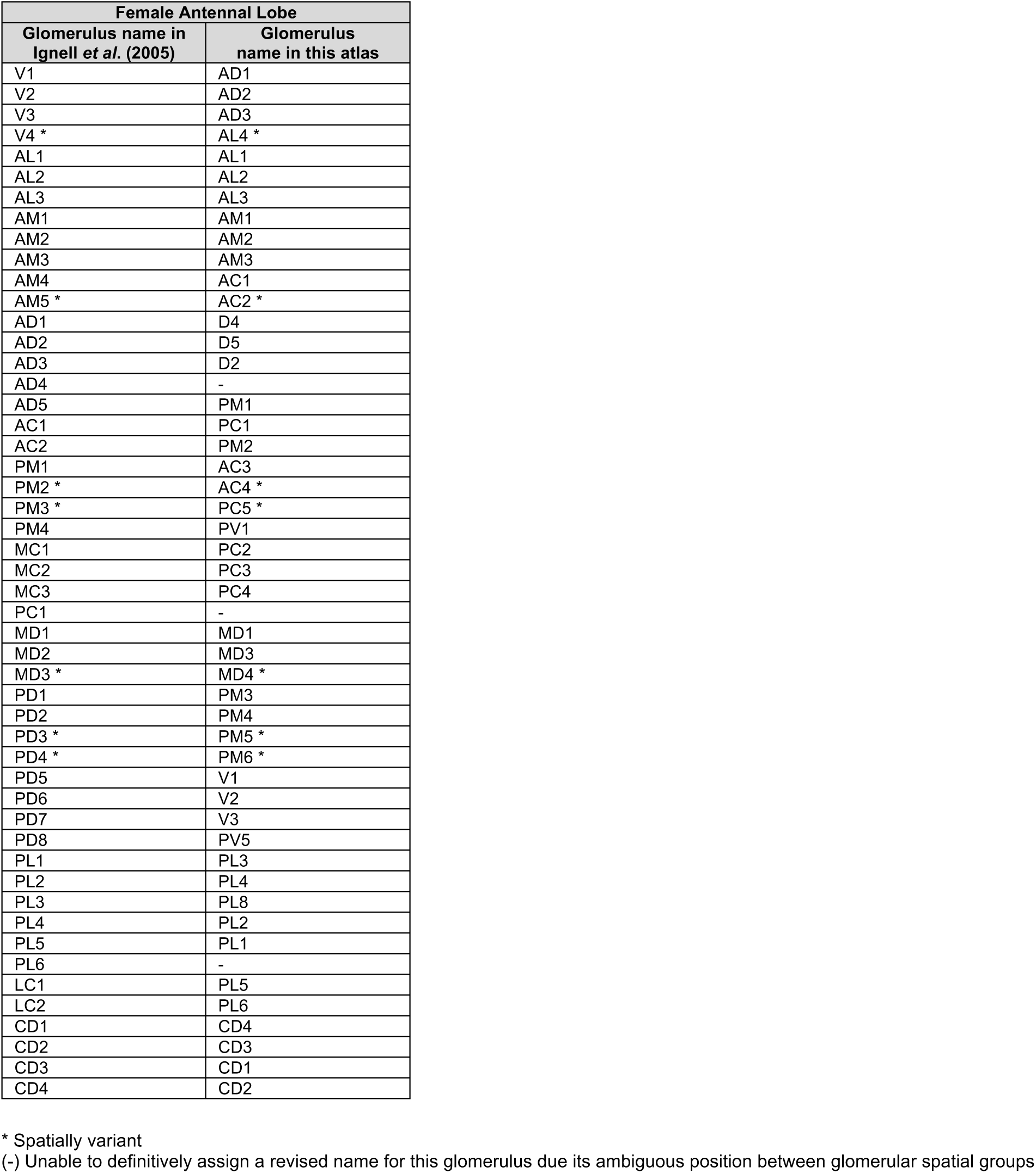
Revised names for all glomeruli annotated within the female antennal lobe model of Ignell *et al*. (2005)

| Female Antennal Lobe |  |
| --- | --- |
| Glomerulus name in Ignell *et al.* (2005) | Glomerulus name in this atlas |
| V1 | AD1 |
| V2 | AD2 |
| V3 | AD3 |
| V4 * | AL4 * |
| AL1 | AL1 |
| AL2 | AL2 |
| AL3 | AL3 |
| AM1 | AM1 |
| AM2 | AM2 |
| AM3 | AM3 |
| AM4 | AC1 |
| AM5 * | AC2 * |
| AD1 | D4 |
| AD2 | D5 |
| AD3 | D2 |
| AD4 | - |
| AD5 | PM1 |
| AC1 | PC1 |
| AC2 | PM2 |
| PM1 | AC3 |
| PM2 * | AC4 * |
| PM3 * | PC5 * |
| PM4 | PV1 |
| MC1 | PC2 |
| MC2 | PC3 |
| MC3 | PC4 |
| PC1 | - |
| MD1 | MD1 |
| MD2 | MD3 |
| MD3 * | MD4 * |
| PD1 | PM3 |
| PD2 | PM4 |
| PD3 * | PM5 * |
| PD4 * | PM6 * |
| PD5 | V1 |
| PD6 | V2 |
| PD7 | V3 |
| PD8 | PV5 |
| PL1 | PL3 |
| PL2 | PL4 |
| PL3 | PL8 |
| PL4 | PL2 |
| PL5 | PL1 |
| PL6 | - |
| LC1 | PL5 |
| LC2 | PL6 |
| CD1 | CD4 |
| CD2 | CD3 |
| CD3 | CD1 |
| CD4 | CD2 |

\* Spatially variant
(-) Unable to definitively assign a revised name for this glomerulus due its ambiguous position between glomerular spatial groups

**Table S4.** Revised names for all glomeruli annotated in the male antennal lobe model of Ignell *et al*. (2005)

| Male Antennal Lobe |  |
| --- | --- |
| Glomerulus name in Ignell *et al.* (2005) | Glomerulus name in this atlas |
| V1 | AD3 |
| V2 | AD1 |
| V3 | AD2 |
| AL1 | AL1 |
| AL2 | AL2 |
| AL3 | AL3 |
| AL4 * | AL4 * |
| AM1 | AM1 |
| AM2 | AM2 |
| AM3 | AM3 |
| AM4 * | AC2 * |
| AM5 | AC3 |
| AM6 | D1 |
| AM7 * | D3 * |
| AD1 | D4 |
| AD2 | D5 |
| AD3 | D2 |
| AD4 | - |
| AD5 | PM1 |
| AD6 | PM2 |
| AD7 | PM3 |
| AC1 | PC1 |
| AC2 | AC1 |
| AC3 * | AD5 * |
| AC4 * | AD4 * |
| PM1 * | AC4 * |
| PM2 | PV1 |
| PM3 * | PM6 * |
| PM4 * | PM5 * |
| PC1 | PC4 |
| PC2 | CD4 |
| PC3 | CD5 |
| MD1 | MD1 |
| MD2 | MD3 |
| MD3 * | MD4 * |
| PD1 | PM4 |
| PD2 | V1 |
| PD3 | V2 |
| PD4 | V4 |
| PD5 | V3 |
| PD6 | V6 |
| PD7 | V7 |
| PD8 | V8 |
| PL1 | PL2 |
| PL2 | PL1 |
| LC1 | PL3 |
| CD1 | CD1 |
| CD2 | CD3 |
| CD3 | CD2 |

\* Spatially variant
(-) Unable to definitively assign a revised name for this glomerulus due its ambiguous position between glomerular spatial groups

**Table S5:** Quantitative comparison of glomerular volume in the female and male antennal lobe.

| Glomerulus | Female |  | *n* | Male |  | *n* | *U*-test for right and left AL of female | *U*-test for right and left AL of male | *U*-test for female and male |
| --- | --- | --- | --- | --- | --- | --- | --- | --- | --- |
|  | Volume (10<sup>3</sup> μm<sup>3</sup>) | Relative volume (%) |  | Volume (10<sup>3</sup> μm<sup>3</sup>) | Relative volume (%) |  | *P*-value | *P*-value | *P*-value |
| AC1 | 1.34 ± 0.46 | 1.07 ± 0.31 | 10 | 1.24 ± 0.52 | 1.48 ± 0.66 | 9 | 0.310 | 1.000 | 0.133 |
| AC3 | 1.24 ± 0.47 | 0.99 ± 0.35 | 10 | 1.05 ± 1.04 | 1.20 ± 1.17 | 8 | 0.841 | 0.486 | 0.460 |
| AL1 | 3.14 ± 1.03 | 2.56 ± 0.83 | 10 | 2.04 ± 1.15 | 2.38 ± 1.31 | 10 | 0.421 | 0.421 | 0.529 |
| AL2 | 2.04 ± 0.73 | 1.66 ± 0.58 | 10 | 1.19 ± 0.72 | 1.38 ± 0.86 | 10 | 0.421 | 1.000 | 0.280 |
| AL3 | 1.73 ± 0.74 | 1.37 ± 0.54 | 10 | 0.78 ± 0.54 | 0.88 ± 0.55 | 9 | 0.841 | 0.730 | 0.065 |
| AM1 | 3.89 ± 1.31 | 3.10 ± 0.78 | 10 | 1.86 ± 1.00 | 2.19 ± 1.24 | 10 | 0.151 | 0.691 | 0.089 |
| AM2 | 2.15 ± 1.07 | 1.73 ± 0.86 | 10 | 1.95 ± 0.75 | 2.29 ± 0.85 | 10 | 0.691 | 0.222 | 0.105 |
| AM3 | 1.31 ± 0.61 | 1.06 ± 0.47 | 10 | 1.43 ± 0.36 | 1.67 ± 0.36 | 10 | 1.000 | 0.421 | 0.004* |
| AD1 | 4.71 ± 0.83 | 3.82 ± 0.56 | 10 | 2.46 ± 0.67 | 2.89 ± 0.76 | 10 | 0.421 | 0.056 | 0.005* |
| AD2 | 1.33 ± 0.68 | 1.09 ± 0.58 | 10 | 0.81 ± 0.64 | 0.93 ± 0.67 | 10 | 1.000 | 0.548 | 0.280 |
| AD3 | 1.03 ± 0.58 | 0.84 ± 0.46 | 10 | 0.76 ± 0.55 | 0.90 ± 0.68 | 9 | 0.222 | 0.111 | 1.000 |
| VC1 | 4.18 ± 1.72 | 3.33 ± 1.30 | 10 | 1.82 ± 0.53 | 2.14 ± 0.64 | 10 | 1.000 | 0.310 | 0.036* |
| VC2 | 2.44 ± 0.98 | 1.95 ± 0.72 | 10 | 1.51 ± 0.68 | 1.75 ± 0.79 | 10 | 0.691 | 0.151 | 0.029 |
| VC3 | 3.39 ± 2.26 | 2.64 ± 1.57 | 10 | 1.31 ± 0.67 | 1.50 ± 0.64 | 10 | 0.548 | 0.691 | 0.123 |
| VC4 | 2.53 ± 2.58 | 1.99 ± 2.05 | 10 | 1.27 ± 0.62 | 1.50 ± 0.76 | 10 | 0.841 | 0.841 | 0.912 |
| VC5 | 2.38 ± 1.55 | 1.88 ± 1.03 | 10 | 1.60 ± 0.62 | 1.87 ± 0.72 | 10 | 0.008* | 0.841 | 0.436 |
| VC6 | 3.13 ± 2.15 | 2.52 ± 1.60 | 10 | 2.77 ± 1.15 | 3.23 ± 1.29 | 10 | 0.841 | 1.000 | 0.315 |
| VC7 | 1.03 ± 0.85 | 0.84 ± 0.67 | 10 | 0.87 ± 0.54 | 1.02 ± 0.61 | 9 | 0.032* | 0.016* | 0.315 |
| VC8 | 1.07 ± 0.69 | 0.87 ± 0.53 | 9 | 0.51 ± 0.26 | 0.60 ± 0.30 | 10 | 0.730 | 0.151 | 0.356 |
| VC9 | 0.73 ± 0.48 | 0.61 ± 0.43 | 9 | 0.47 ± 0.27 | 0.55 ± 0.31 | 9 | 0.730 | 0.191 | 0.730 |
| PC1 | 0.55 ± 0.41 | 0.46 ± 0.34 | 10 | 0.40 ± 0.32 | 0.47 ± 0.38 | 10 | 0.032 | 0.421 | 0.912 |
| PC2 | 0.66 ± 0.37 | 0.55 ± 0.30 | 9 | 0.68 ± 0.60 | 0.78 ± 0.68 | 10 | 0.064 | 0.691 | 0.720 |
| PC3 | 0.65 ± 0.36 | 0.53 ± 0.31 | 8 | 0.40 ± 0.20 | 0.46 ± 0.23 | 9 | 0.886 | 0.905 | 0.673 |
| PC4 | 0.64 ± 0.41 | 0.53 ± 0.37 | 10 | 0.48 ± 0.26 | 0.55 ± 0.28 | 9 | 0.056 | 0.111 | 0.661 |
| PV1 | 2.30 ± 0.71 | 1.86 ± 0.51 | 10 | 1.98 ± 0.59 | 2.35 ± 0.76 | 10 | 0.548 | 0.056 | 0.166 |
| PV2 | 2.01 ± 1.21 | 1.61 ± 0.90 | 10 | 1.91 ± 1.44 | 2.20 ± 1.60 | 10 | 0.056 | 1.000 | 0.481 |
| PV3 | 2.24 ± 1.43 | 1.80 ± 1.12 | 10 | 1.81 ± 1.25 | 2.07 ± 1.41 | 10 | 0.841 | 0.151 | 0.912 |
| PV4 | 0.98 ± 0.51 | 0.79 ± 0.40 | 10 | 0.68 ± 0.38 | 0.79 ± 0.43 | 10 | 0.151 | 0.310 | 0.796 |
| PV5 | 0.75 ± 0.62 | 0.59 ± 0.44 | 10 | 0.61 ± 0.25 | 0.71 ± 0.27 | 10 | 0.841 | 0.691 | 0.248 |
| PM1 | 0.64 ± 0.44 | 0.53 ± 0.36 | 10 | 0.60 ± 0.27 | 0.69 ± 0.30 | 9 | 0.222 | 0.191 | 0.243 |
| PM2 | 0.74 ± 0.45 | 0.61 ± 0.37 | 9 | 0.59 ± 0.27 | 0.68 ± 0.35 | 9 | 0.413 | 0.730 | 0.667 |
| PM3 | 0.88 ± 0.53 | 0.73 ± 0.45 | 9 | 0.53 ± 0.44 | 0.63 ± 0.57 | 10 | 0.413 | 0.222 | 0.400 |
| PM4 | 0.92 ± 0.46 | 0.77 ± 0.42 | 9 | 0.49 ± 0.30 | 0.58 ± 0.35 | 9 | 0.556 | 1.000 | 0.297 |
| PL1 | 2.14 ± 1.32 | 1.77 ± 1.09 | 10 | 1.61 ± 1.04 | 1.85 ± 1.10 | 10 | 1.000 | 0.056 | 0.971 |
| PL2 | 1.70 ± 1.07 | 1.40 ± 0.87 | 10 | 1.75 ± 1.29 | 2.05 ± 1.54 | 10 | 0.008* | 0.095 | 0.481 |
| PL3 | 2.95 ± 1.26 | 2.44 ± 1.07 | 10 | 1.70 ± 0.86 | 2.04 ± 1.12 | 10 | 1.000 | 0.548 | 0.393 |
| PL4 | 2.10 ± 1.12 | 1.74 ± 0.92 | 10 | 1.22 ± 1.16 | 1.46 ± 1.41 | 10 | 0.421 | 0.421 | 0.248 |
| PL5 | 1.21 ± 0.87 | 0.97 ± 0.62 | 10 | 1.06 ± 0.63 | 1.25 ± 0.76 | 10 | 0.421 | 0.222 | 0.481 |
| PL6 | 1.05 ± 0.62 | 0.86 ± 0.55 | 10 | 1.29 ± 1.28 | 1.49 ± 1.39 | 10 | 0.151 | 1.000 | 0.166 |
| PL7 | 0.96 ± 0.68 | 0.80 ± 0.61 | 10 | 0.81 ± 0.66 | 0.99 ± 0.80 | 8 | 0.310 | 0.686 | 0.829 |
| PL8 | 1.86 ± 1.23 | 1.54 ± 1.08 | 10 | 0.85 ± 0.49 | 1.00 ± 0.57 | 9 | 0.548 | 0.905 | 0.243 |
| PL9 | 2.40 ± 0.73 | 1.97 ± 0.68 | 9 | 2.23 ± 1.40 | 2.64 ± 1.59 | 9 | 1.000 | 0.413 | 0.340 |
| D1 | 1.79 ± 1.74 | 1.42 ± 1.36 | 10 | 1.04 ± 0.39 | 1.24 ± 0.51 | 10 | 0.032* | 0.222 | 0.529 |
| D2 | 1.06 ± 0.78 | 0.86 ± 0.62 | 10 | 0.63 ± 0.42 | 0.76 ± 0.55 | 10 | 0.841 | 0.691 | 0.481 |
| D4 | 1.21 ± 0.54 | 1.01 ± 0.47 | 9 | 1.39 ± 0.67 | 1.66 ± 0.86 | 10 | 0.905 | 0.421 | 0.113 |
| D5 | 1.73 ± 1.33 | 1.47 ± 1.17 | 10 | 1.14 ± 0.64 | 1.31 ± 0.67 | 9 | 0.841 | 0.730 | 0.968 |
| CD1 | 0.80 ± 0.41 | 0.64 ± 0.32 | 10 | 0.76 ± 0.53 | 0.91 ± 0.65 | 10 | 0.691 | 0.841 | 0.393 |
| CD2 | 0.87 ± 0.51 | 0.72 ± 0.45 | 10 | 0.60 ± 0.43 | 0.73 ± 0.55 | 10 | 0.548 | 0.691 | 0.853 |
| CD3 | 0.59 ± 0.32 | 0.50 ± 0.29 | 10 | 0.78 ± 0.30 | 0.93 ± 0.39 | 10 | 0.548 | 1.000 | 0.019* |
| CD4 | 1.19 ± 0.49 | 1.00 ± 0.49 | 10 | 1.52 ± 0.58 | 1.77 ± 0.64 | 10 | 0.151 | 1.000 | 0.015* |
| CD5 | 0.97 ± 0.73 | 0.80 ± 0.62 | 10 | 0.80 ± 0.50 | 0.92 ± 0.55 | 10 | 0.841 | 0.222 | 0.684 |
| CD6 | 0.80 ± 0.62 | 0.66 ± 0.50 | 9 | 0.53 ± 0.19 | 0.63 ± 0.22 | 10 | 0.064 | 0.841 | 0.661 |
| V1 | 2.29 ± 1.46 | 1.90 ± 1.28 | 10 | 1.68 ± 0.79 | 1.98 ± 0.99 | 10 | 0.421 | 0.841 | 0.579 |
| V2 | 1.51 ± 0.90 | 1.20 ± 0.68 | 10 | 0.82 ± 0.38 | 0.96 ± 0.41 | 10 | 0.008* | 0.095 | 0.579 |
| V3 | 1.69 ± 0.95 | 1.33 ± 0.69 | 10 | 1.12 ± 1.02 | 1.25 ± 1.04 | 10 | 0.691 | 0.691 | 0.796 |
| V4 | 1.38 ± 0.93 | 1.14 ± 0.80 | 10 | 0.62 ± 0.36 | 0.74 ± 0.42 | 10 | 0.421 | 0.691 | 0.315 |
| V5 | 1.15 ± 0.52 | 0.94 ± 0.42 | 10 | 0.88 ± 0.57 | 1.02 ± 0.62 | 10 | 0.691 | 0.691 | 0.971 |
| V6 | 1.04 ± 0.93 | 0.86 ± 0.80 | 10 | 1.23 ± 1.13 | 1.43 ± 1.30 | 10 | 0.691 | 0.222 | 0.143 |
| V7 | 0.74 ± 0.41 | 0.60 ± 0.34 | 10 | 1.10 ± 0.96 | 1.26 ± 1.08 | 9 | 0.548 | 0.556 | 0.400 |
| V8 | 0.94 ± 0.40 | 0.76 ± 0.31 | 9 | 0.96 ± 0.56 | 1.11 ± 0.60 | 10 | 1.000 | 0.691 | 0.182 |
| MD1 | 5.22 ± 1.09 | 4.26 ± 0.94 | 10 | 4.09 ± 0.63 | 4.81 ± 0.81 | 10 | 0.691 | 0.032* | 0.218 |
| MD2 | 1.84 ± 1.03 | 1.52 ± 0.80 | 9 | 1.27 ± 0.64 | 1.48 ± 0.71 | 10 | 0.191 | 1.000 | 0.720 |
| MD3 | 1.33 ± 0.44 | 1.10 ± 0.35 | 8 | 1.44 ± 0.61 | 1.70 ± 0.69 | 10 | 0.114 | 1.000 | 0.016* |
| Total AL Volume | 123.32 ± 12.03 |  | 10 | 85.37 ± 7.65 |  | 10 | 0.548 | 0.548 | <0.001* |

Volume and relative volume are presented as Mean ± SD. Relative volume was used for statistical comparison to control for variation in the absolute volume in individuals, except for comparing total antennal lobe volumes. Asterisks indicate statistical significance at the level of *P* = 0.05.

**Figure S1.**
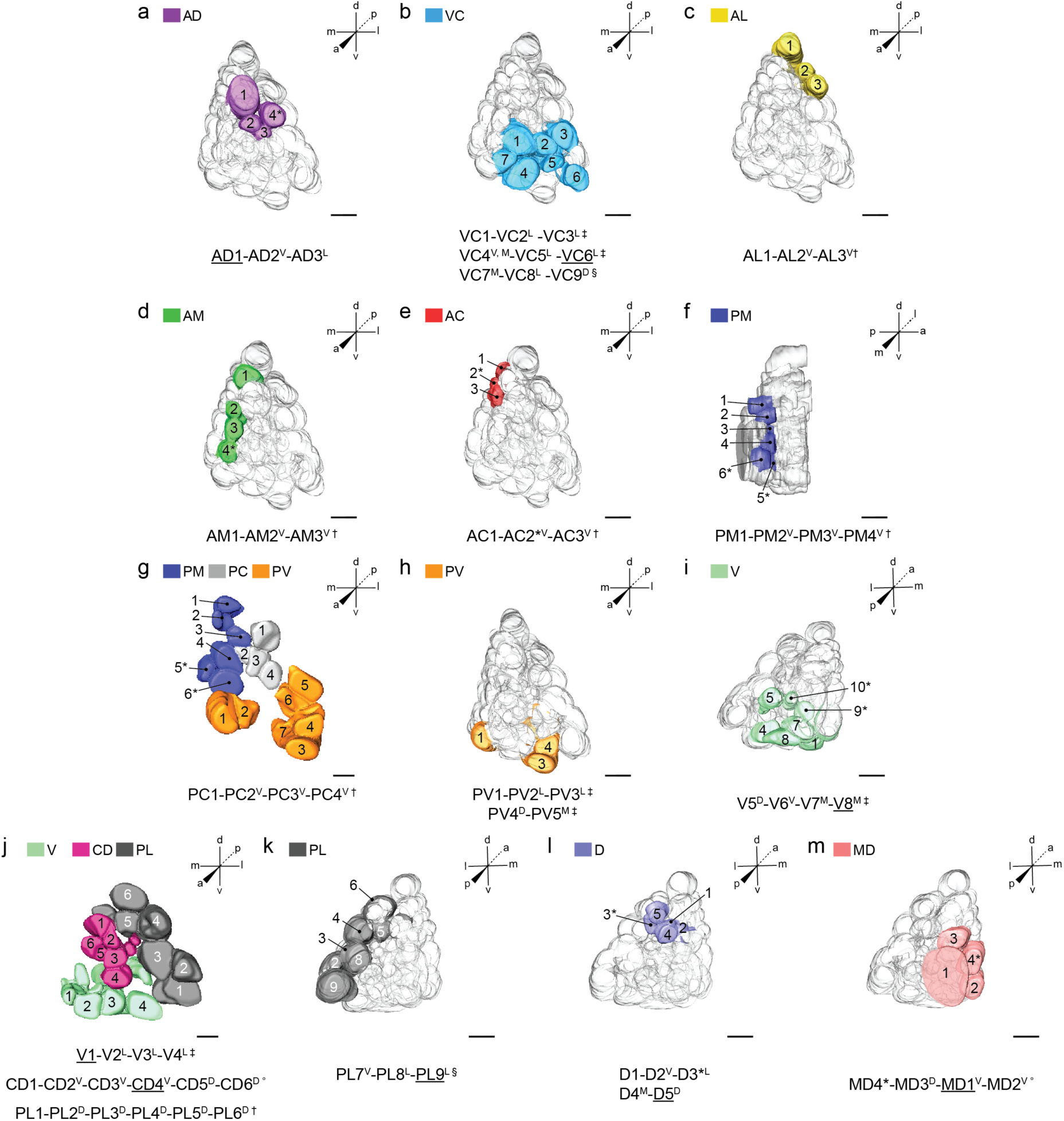
Intra-group glomerular arrays highlighted on an example 3D model of a left antennal lobe from a nc82-stained LVPib12 female brain. (**a – m**) Glomeruli within intra-group arrays belonging to the thirteen color-coded spatial groups are highlighted from various perspectives of the antennal lobe. Scale bars = 10 µm. All remaining glomeruli were made transparent to highlight these example arrays and their typical spatial position. Textual annotations that denote the typical spatial arrangement of glomeruli within intra-group glomerular arrays are shown below each perspective of the model lobe. Underlined glomeruli in arrays (AD1, VC6, D5, V1, V8, MD1, CD4 and PL9) are landmarks that facilitate identification of other glomeruli based on their relative spatial position. Glomeruli are typically arrayed in columns^†^, rows^‡^, circles^°^, or randomly arrayed and numbered based on depth^§^. Each glomerulus may occupy a medial^M^, lateral^L^, dorsal^D^ or ventral^V^ position relative to the preceding glomerulus in its array. Landmark glomeruli are underlined. Commonly observed variant glomeruli are indicated by an asterisk*. A full series of intra- and inter-group glomerular arrays are detailed in Table 1.

**Figure S2.**
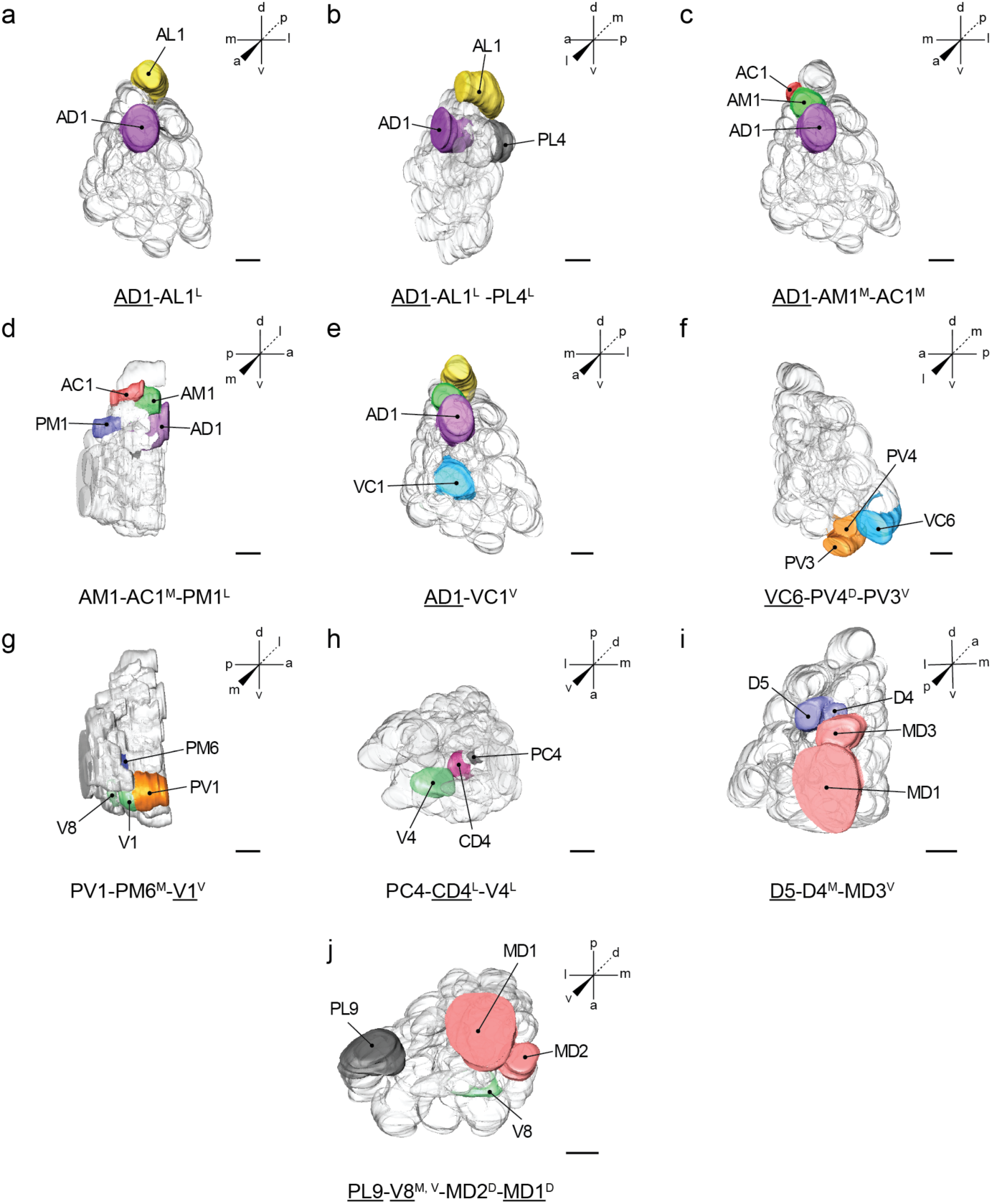
Inter-group glomerular arrays highlighted on an example 3D model of a left antennal lobe from a nc82-stained LVPib12 female brain. (**a – j**) Glomeruli within inter-group arrays are highlighted from various perspectives of the antennal lobe and are color-coded on the 3D model according to their spatial group. Scale bars = 10 µm. All remaining glomeruli were made transparent to highlight these example arrays and their typical spatial position. Glomeruli are arranged in layers from the anterior through posterior sections of the antennal lobe. Textual annotations that denote the typical spatial arrangement of glomeruli within an inter-group glomerular array are shown below each perspective of the model lobe. Underlined glomeruli in arrays (AD1, VC6, D5, V1, V8, MD1, CD4 and PL9) are landmarks that facilitate identification of other glomeruli based on their relative spatial position. Each glomerulus may occupy a medial^M^, lateral^L^, dorsal^D^ or ventral^V^ position relative to the preceding glomerulus in its array. A full series of intra- and inter-group glomerular arrays are detailed in Table 1.

**Figure S3.**
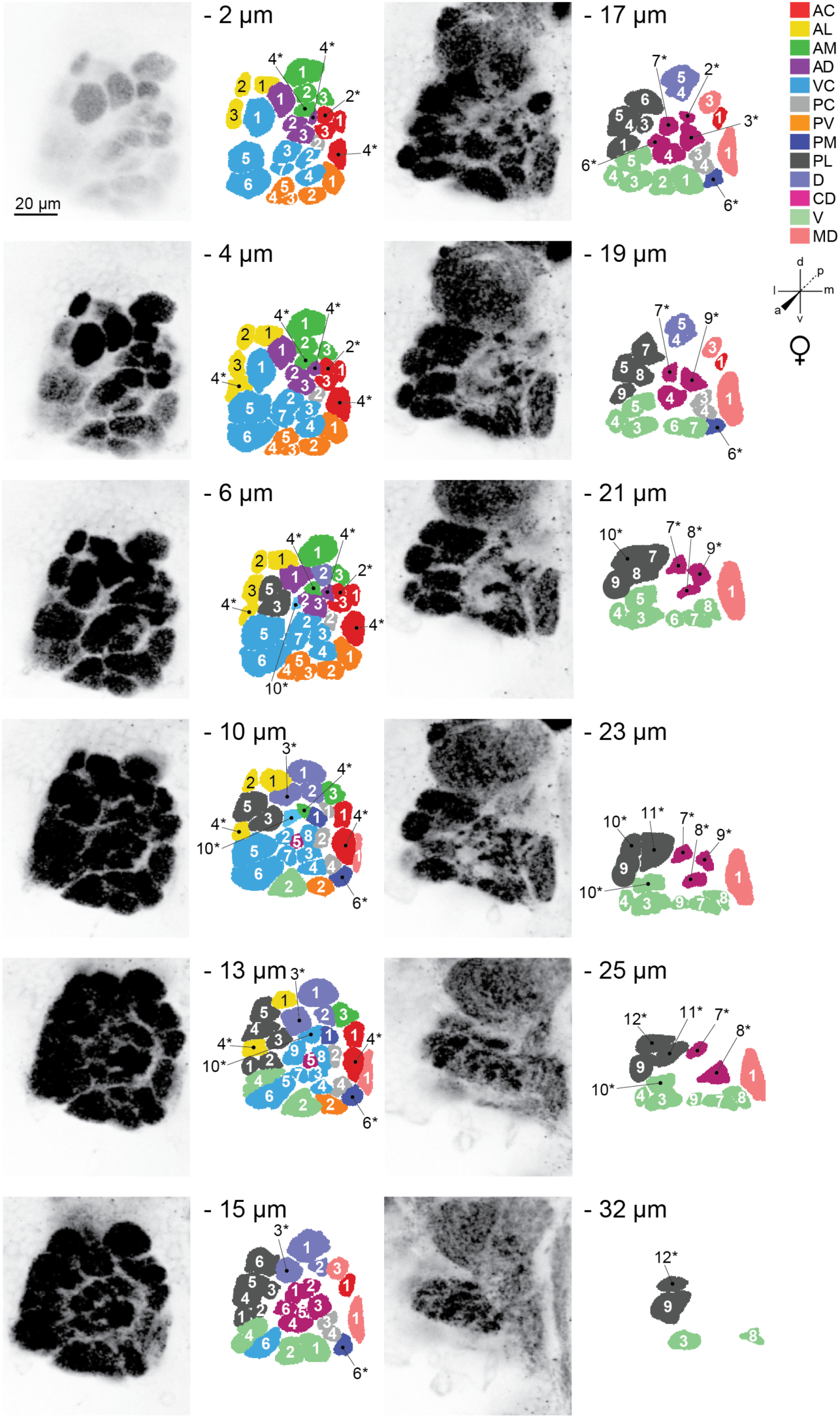
Representative confocal stack from the right antennal lobe of an adult female LVPib12 *Aedes aegypti* as shown by nc82 staining. Twelve frontal planes from a total of 35 images taken at 1µm intervals were selected for illustration of the typical geometric arrangement of glomeruli; Scale bar, 20 µm. The depth of each confocal slice is indicated. Glomeruli within each reconstructed slice are color-coded according to their predicted spatial group. Glomeruli are numbered, with 61 out of 63 spatially invariant glomeruli evident in the twelve antennal lobe slices depicted here. Variant glomeruli are indicated by asterisks.

**Figure S4.**
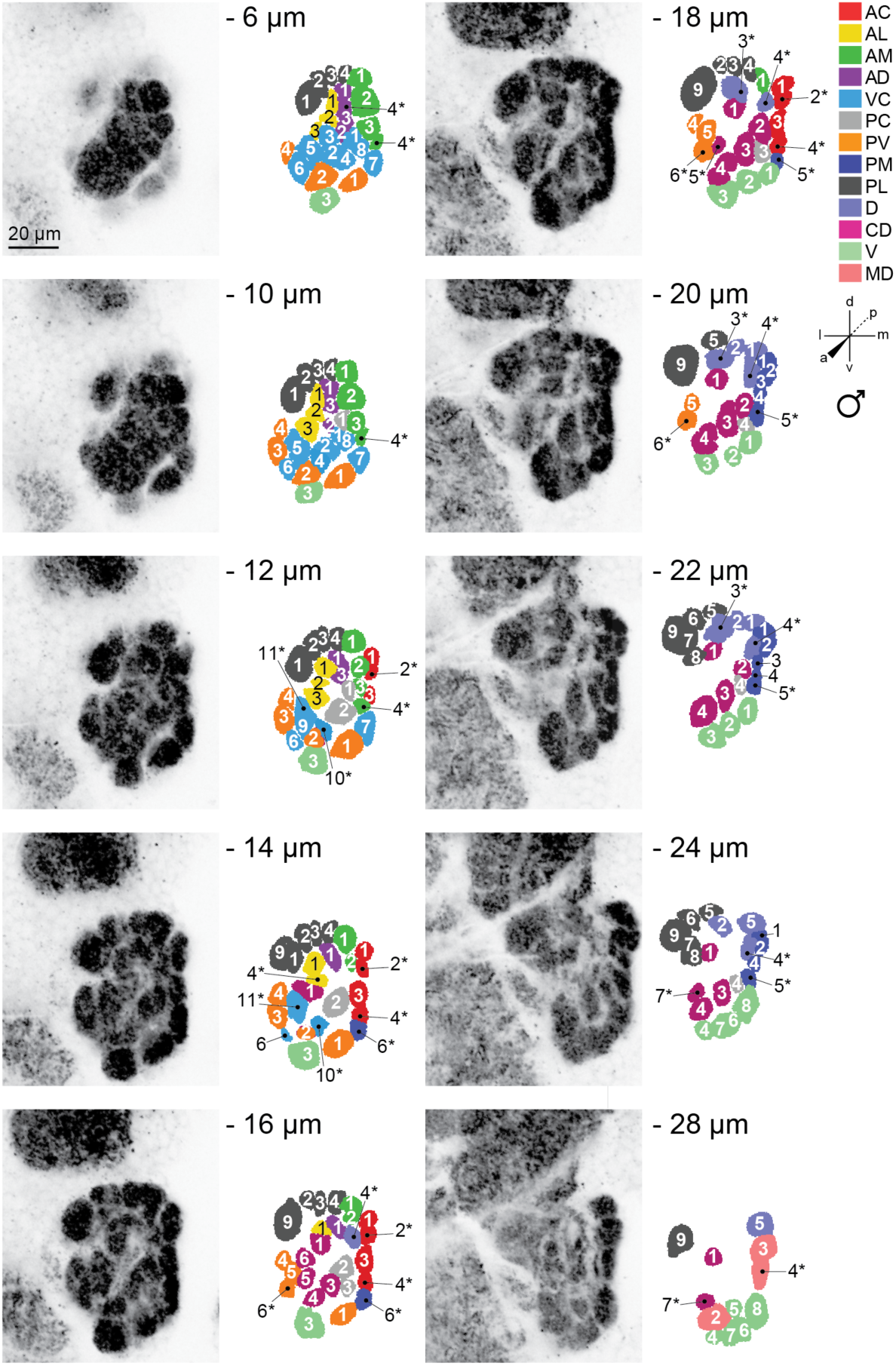
Representative confocal stack from the right antennal lobe of an adult male LVPib12 *Aedes aegypti* as shown by nc82 staining. Ten frontal planes from a total of 40 images taken at 1µm intervals were selected for illustration of the typical geometric arrangement of glomeruli; Scale bar, 20 µm. The depth of each confocal slice is indicated. Glomeruli within each reconstructed slice are color-coded according to their predicted spatial group. Glomeruli are numbered, with 63 out of 63 spatially invariant glomeruli evident in the ten antennal lobe slices depicted here. Variant glomeruli are indicated by asterisks.

**Figure S5.**
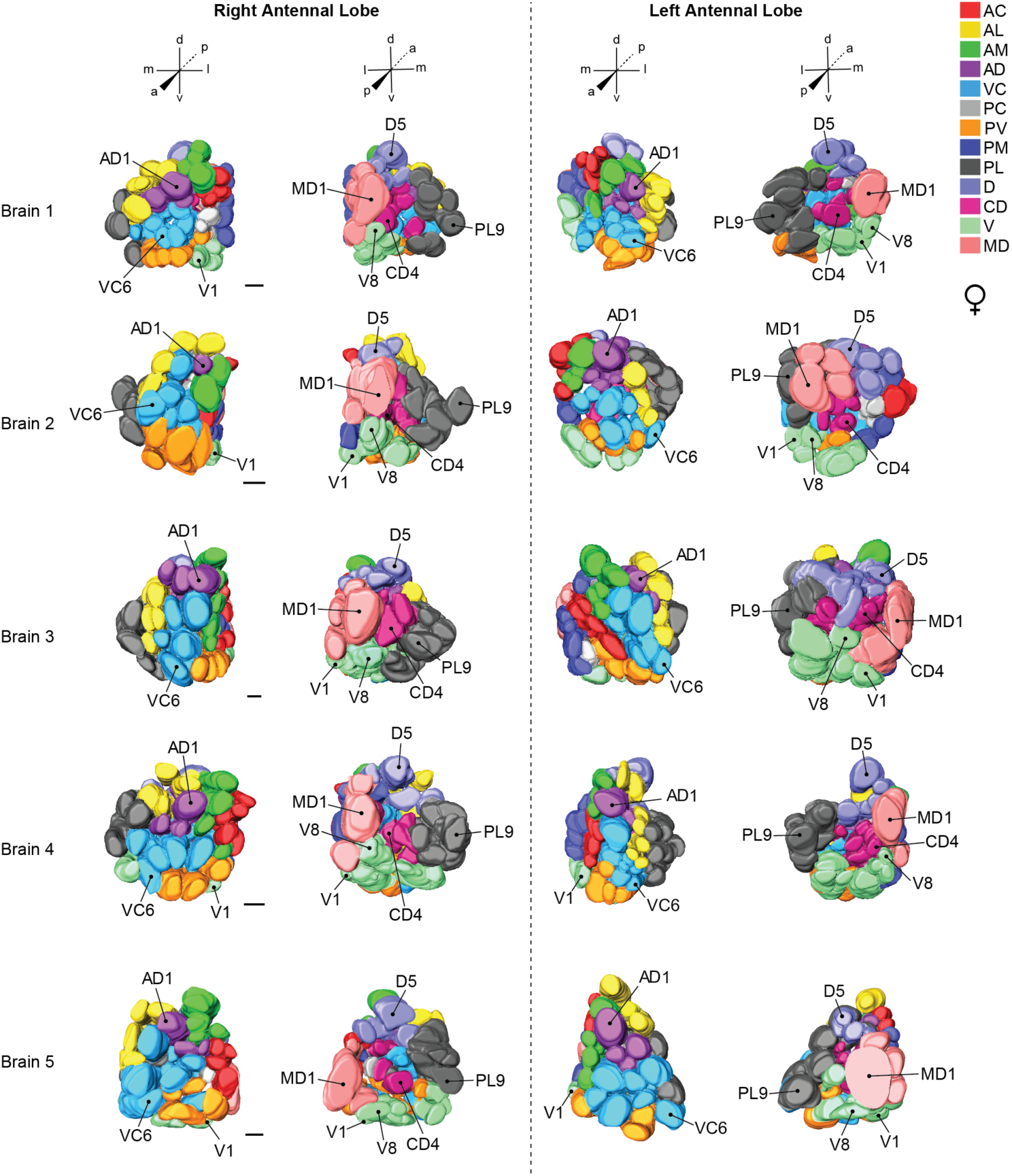
Representative variation in antennal lobe shape in reconstructions from *in vitro* fixed, dissected and stained brains. Three dimensional models of the right and left antennal lobes from 5 brain samples of LVPib12 females stained with nc82 antibody are shown. Anterior and posterior perspectives of each antennal lobe reconstruction are illustrated. Glomeruli are color-coded in the 3D models according to their spatial group. Landmark glomeruli are labelled on all reconstructed antennal lobes and include AD1, VC6 located on the anterior surface and D5, V1, V8, MD1, CD4 and PL9 on the posterior surface of the antennal lobe. Scale bars = 10 µm.

**Figure S6.**
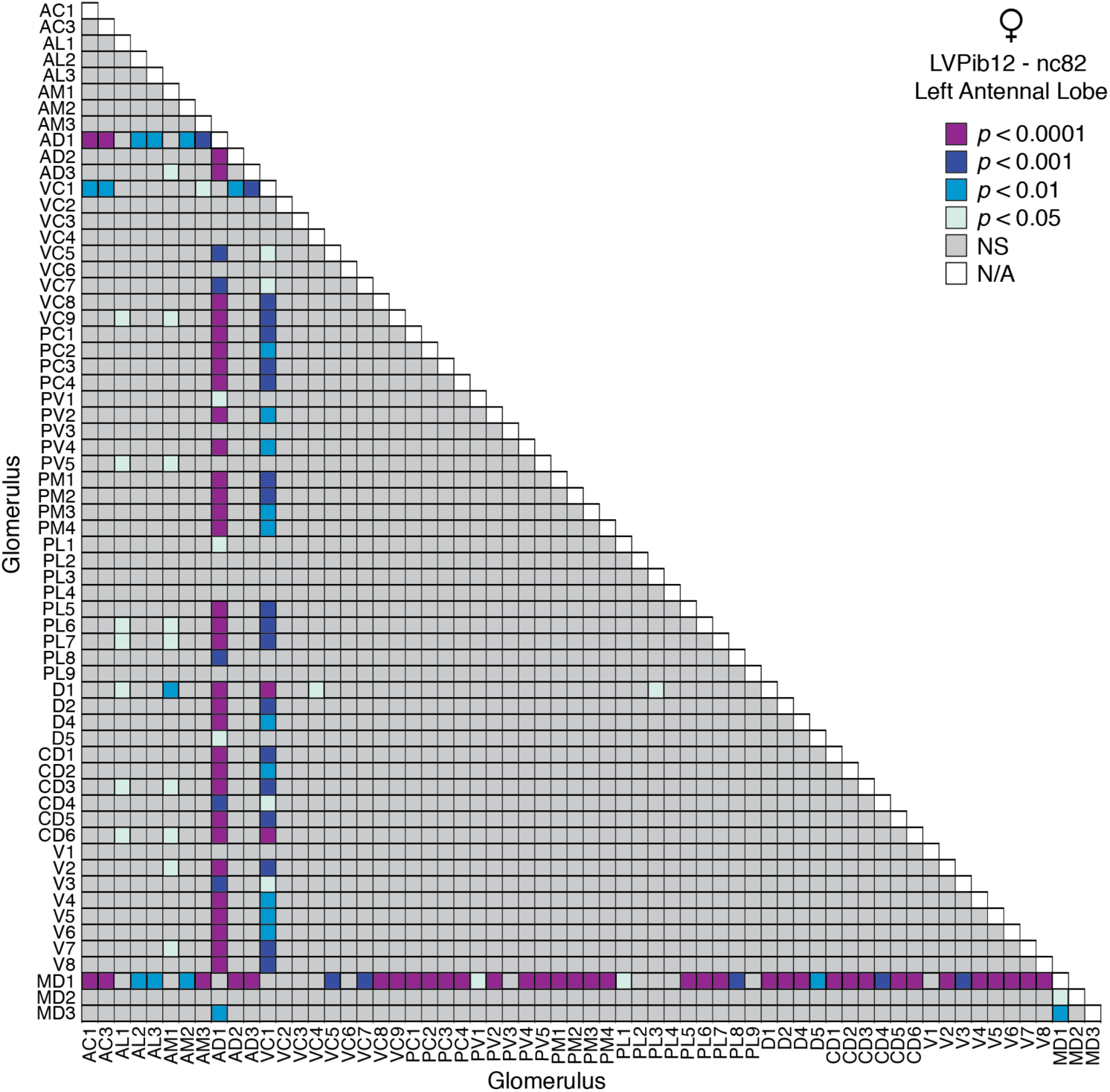
Probability heatmap visually representing the statistical significance of pairwise post-hoc Tukey’s HSD tests comparing the mean volumes of all 63 spatially invariant glomeruli in left antennal lobe of nc82-stained brains from adult female LVPib12 *Aedes aegypti* (n = 5 brains). Multiplicity adjusted *P* values are plotted for each comparison. AD1, VC1 and MD1 typically had larger volumes when compared with all other glomeruli. Glomerular means in this lobe differed significantly as determined by one-way ANOVA: F (62, 247) = 5.412, *P*<0.0001. Abbreviations: NS (not significant). N/A (comparisons between volumetric means of the same glomerulus are not applicable).

**Figure S7.**
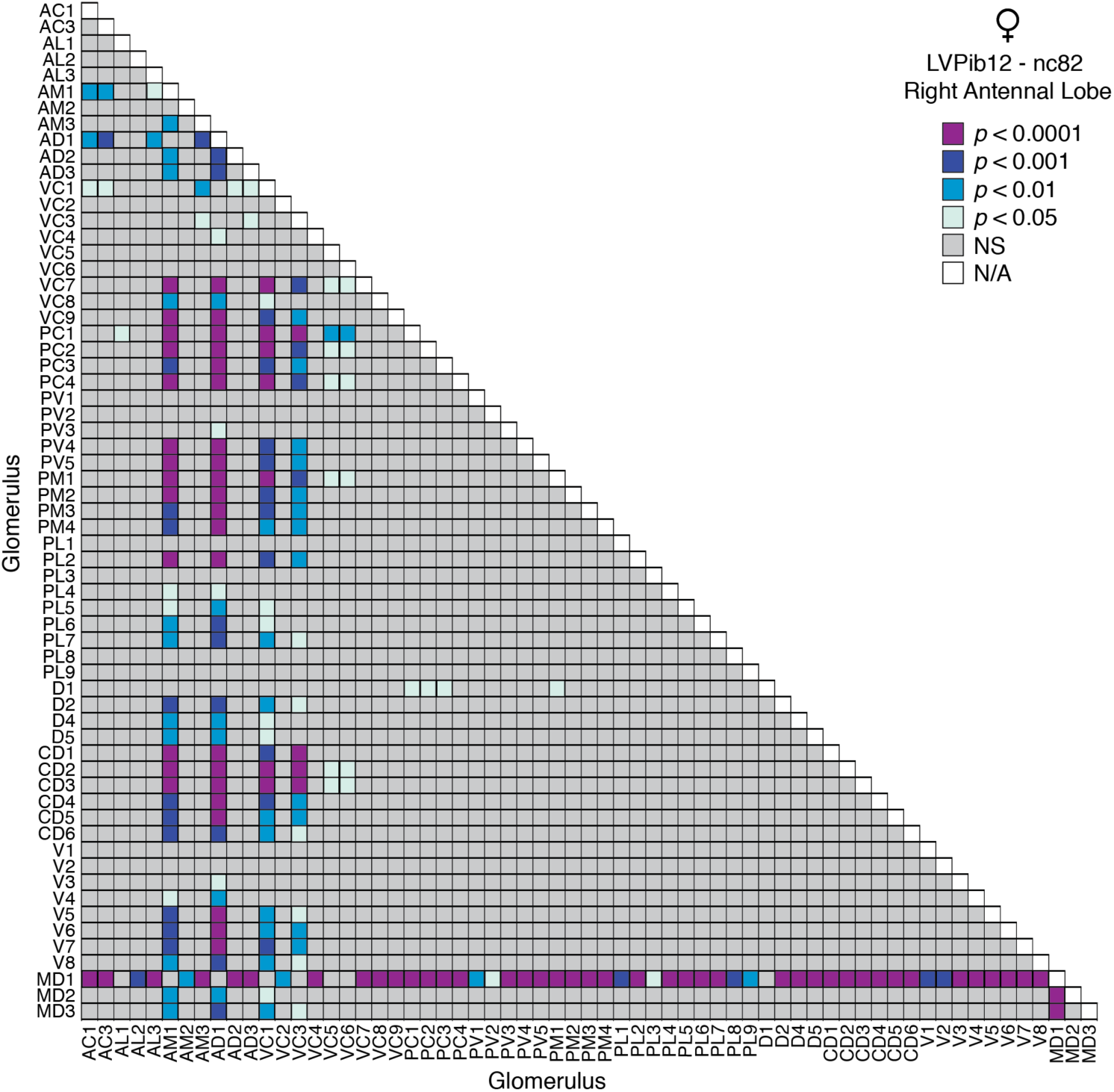
Probability heatmap visually representing the statistical significance of pairwise post-hoc Tukey’s HSD tests comparing the mean volumes of all 63 spatially invariant glomeruli in right antennal lobe of nc82-stained brains from adult female LVPib12 *Aedes aegypti* (n = 5 brains). Multiplicity adjusted *P* values are plotted for each comparison. AM1, AD1, VC1, VC3 and MD1 typically had larger volumes when compared with all other glomeruli. Glomerular means in this lobe differed significantly as determined by one-way ANOVA: F (62, 242) = 6.521, *P*<0.0001. Abbreviations: NS (not significant). N/A (comparisons between volumetric means of the same glomerulus are not applicable).

**Figure S8.**
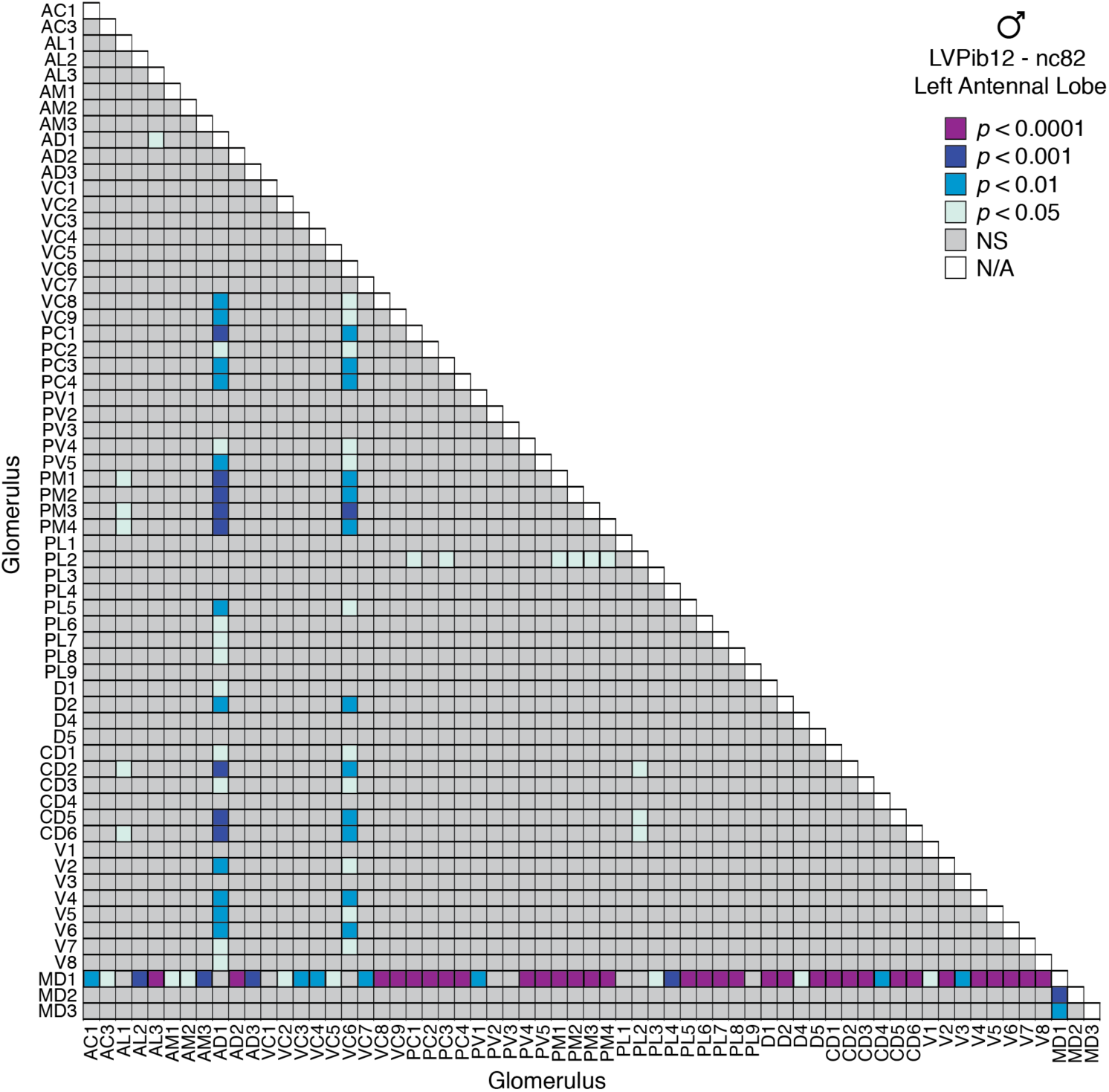
Probability heatmap visually representing the statistical significance of pairwise post-hoc Tukey’s HSD tests comparing the mean volumes of all 63 spatially invariant glomeruli in left antennal lobe of nc82-stained brains from adult male LVPib12 *Aedes aegypti* (n = 5 brains). Multiplicity adjusted *P* values are plotted for each comparison. AD1, VC6 and MD1 typically had larger volumes when compared with all other glomeruli. Glomerular means in this lobe differed significantly as determined by one-way ANOVA: F (62, 241) = 4.472, *P*<0.0001. Abbreviations: NS (not significant). N/A (comparisons between volumetric means of the same glomerulus are not applicable).

**Figure S9.**
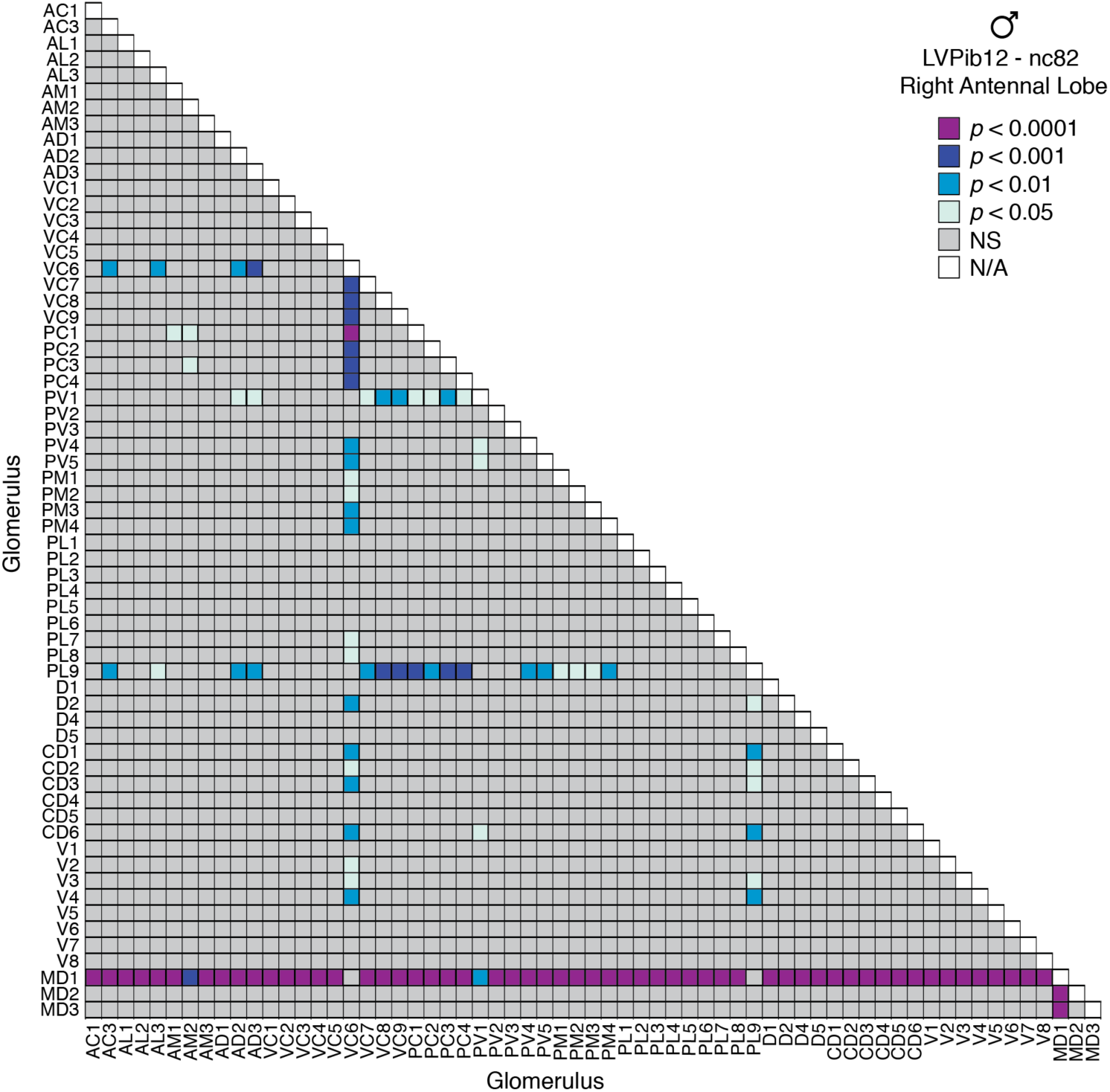
Probability heatmap visually representing the statistical significance of pairwise post-hoc Tukey’s HSD tests comparing the mean volumes of all 63 spatially invariant glomeruli in right antennal lobe of nc82-stained brains from adult male LVPib12 *Aedes aegypti* (n = 5 brains). Multiplicity adjusted *P* values are plotted for each comparison. VC6 and MD1 typically had larger volumes when compared with all other glomeruli. Glomerular means in this lobe differed significantly as determined by one-way ANOVA: F (62, 245) = 5.404, *P*<0.0001. Abbreviations: NS (not significant). N/A (comparisons between volumetric means of the same glomerulus are not applicable).

**Figure S10.**
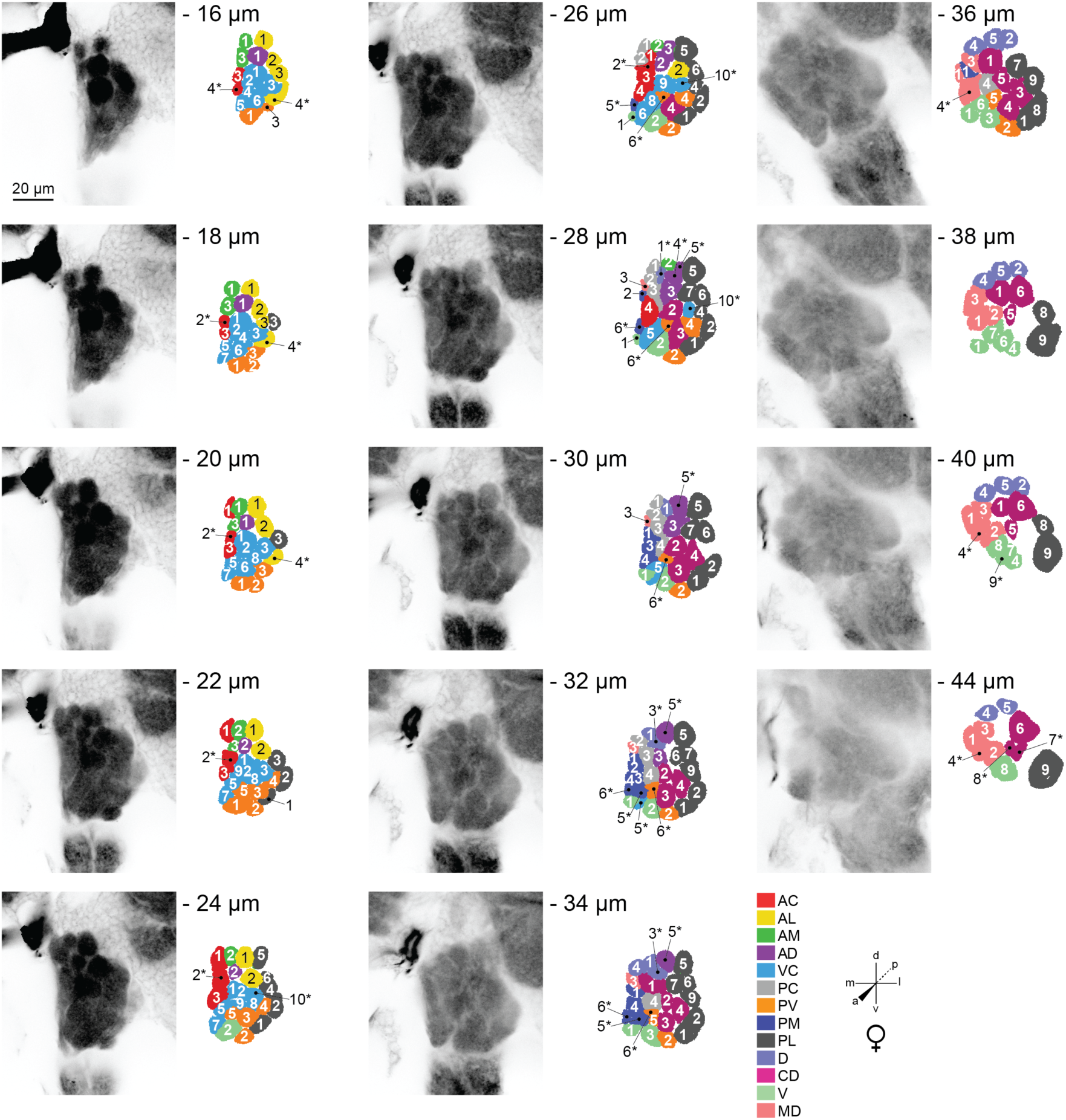
Representative confocal stack from the left antennal lobe of an adult female LVPib12 *Aedes aegypti* as shown by phalloidin staining. Fourteen frontal planes from a total of 45 images taken at 1µm intervals were selected for illustration of the typical geometric arrangement of glomeruli; Scale bar, 20 µm. The depth of each confocal slice is indicated. Glomeruli within each reconstructed slice are color-coded according to their predicted spatial group. Glomeruli are numbered, with 63 out of the 63 spatially invariant glomeruli evident in the fourteen antennal lobe slices depicted here. Variant glomeruli are indicated by asterisks.

**Figure S11.**
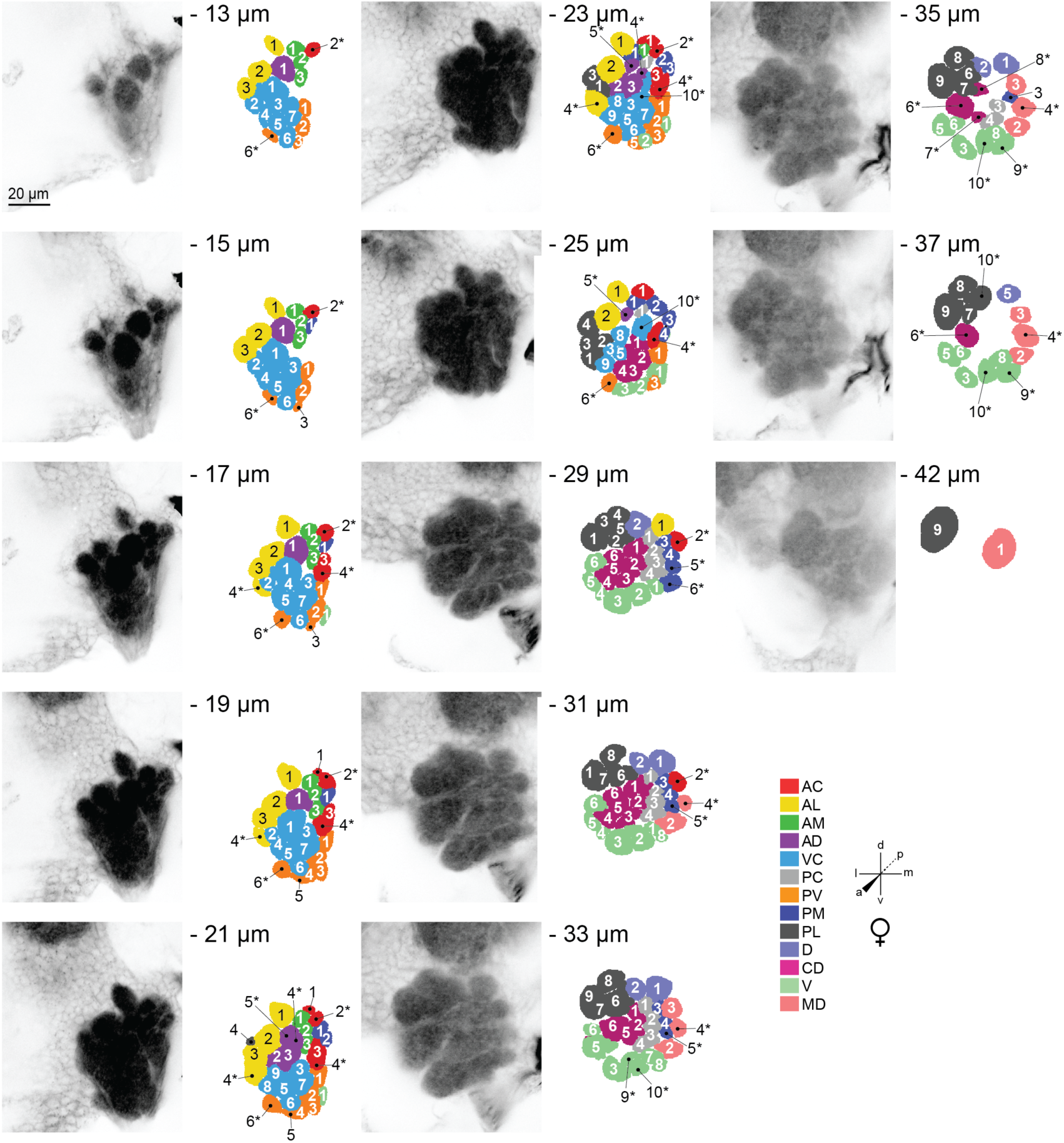
Representative confocal stack from the right antennal lobe of an adult female LVPib12 *Aedes aegypti* as shown by phalloidin staining. Thirteen frontal planes from a total of 45 images taken at 1µm intervals were selected for illustration of the typical geometric arrangement of glomeruli; Scale bar, 20 µm. The depth of each confocal slice is indicated. Glomeruli within each reconstructed slice are color-coded according to their predicted spatial group. Glomeruli are numbered, with 63 out of the 63 spatially invariant glomeruli evident in the thirteen antennal lobe slices depicted here. Variant glomeruli are indicated by asterisks.

